# C1q and immunoglobulins mediate activity-dependent synapse loss in the adult brain

**DOI:** 10.1101/2024.12.18.629085

**Authors:** Gerard Crowley, Emir Turkes, Minjung Kim, Sebastiaan De Schepper, Benjy J. Y. Tan, Javier Rueda-Carrasco, Margarita Toneva, John Christian Fajardo, Judy Z. Ge, Zhengyue Grace Yang, Camille Paoletti, Tammie Sow, David Posner, Annerieke Sierksma, Laís S. S. Ferreira, Dimitra Sokolova, Viktoras Konstantellos, Andrew F. MacAskill, Menna R. Clatworthy, Soyon Hong

## Abstract

C1q, the initiating protein of the classical complement cascade, mediates synapse loss in development and disease. In various mouse models of neurologic diseases, including Alzheimer’s disease, C1q, which is secreted by microglia, the brain’s resident macrophages, is found deposited on synapses in vulnerable brain regions. However, what underlies C1q deposition on synapses in the adult brain is unclear. Using *in vivo* chemogenetics, we demonstrate that neuronal hyperactivity acts as a trigger for region-specific deposition of C1q, which is required for activity-dependent synapse loss. Further, using spatial transcriptomics, live cell tracking, super-resolution microscopy and other molecular and cellular tools, we report a role for B lymphocyte lineage cells and immunoglobulins in the activity-dependent C1q deposition and synapse loss. Overall, our work suggests a link between neuronal hyperactivity and C1q-mediated synapse loss in the adult brain and introduces immunoglobulins as players in this process.

## Main text

### Introduction

Emerging studies suggest that microglia, the brain’s immune cells, regulate synaptic development and function across the lifespan (*1*). During CNS development, neuronal activity regulates synapse pruning (*2–4*) and microglia-mediated elimination of synapses (*5*, *6*). However, whether neuronal activity also underlies synapse loss in diseased adult brains is unclear. Interestingly, region-specific neuronal hyperactivity occurs early in Alzheimer’s disease (AD) patients and mouse models, coinciding with synapse loss (*7*, *8*). However, whether neuronal activity underlies region-specific C1q deposition and synapse loss in adult brains is not known.

Here, we tested the relationship between neuronal activity and C1q deposition using *in vivo* chemogenetics to modulate activity of the perforant pathway, a neuronal circuit that exhibits early dysregulation in AD (*9*). Repeated chemogenetic activation of medial entorhinal cortex (MEC) neurons in wild-type C57Bl6/J (hereafter WT) mice resulted in region-specific C1q deposition and selective synapse elimination in the hippocampus, the primary target of perforant pathway projections. Activity-mediated synapse loss did not occur in *C1qa* knockout mice, suggesting C1q is required for this process. The reverse chemogenetic experiment inhibiting activity along the perforant pathway in the hAPP-J20 (hereafter J20) mouse model of AD, (*10*) reduced C1q deposition and ameliorated synapse loss. Further, we show relevance for immunoglobulins in hyperactivity-induced C1q deposition and synapse loss in WT mice. Altogether, our study suggests a functional crosstalk between neuronal activity, immunoglobulins and C1q-mediated synapse loss in the adult brain.

#### Changes in neuronal activity influence C1q deposition at synapses

First, we sought to determine whether increasing neuronal activity is sufficient to increase C1q deposition at the synapse. Specifically, we targeted the perforant pathway, a neuronal projection connecting MEC to the hippocampal middle part of the dentate gyrus molecular layer (DGML; fig. S1-2). We applied *in vivo* chemogenetics using AAV8-CaMKIIa-HA-hM3Dq-IRES-mCitrine, an excitatory DREADD which is activated upon CNO administration (*11*), and unilaterally targeted the MEC layer II using stereotaxic injection in 3-month-old WT mice. Targeted GFP-immunolabeled projection occurred ipsilaterally from MEC layer II to DGML, allowing for a pairwise comparison with the untargeted contralateral hemisphere from the same mouse brain (fig. S1C) (*12*). It is important to note that the hippocampal regions analyzed are located at least 3.5 mm away from the DREADD injection site, reducing the likelihood of artifacts from surgical procedures. Further, appropriate controls were implemented to reduce off-target effects. For the DREADD virus, separate cohorts of mice were injected with AAV8-CaMKIIa-EGFP as a control. To control for CNO administration, mice injected with either the DREADD or control virus received intraperitoneal injections of saline or CNO once daily for five consecutive days at 7-8 weeks post-surgery and were sacrificed 90 minutes after the final injection (Fig. 1A). CNO treatment in hM3Dq-expressing mice induced an increase in c-Fos levels in the ipsilateral MEC compared to the contralateral MEC (Fig. 1B), suggesting elevated MEC activity. In contrast, no differences in c-Fos levels were observed between the ipsilateral and contralateral MEC in the three control groups: hM3Dq-expressing mice treated with saline and EGFP control virus-expressing mice treated with either CNO or saline (Fig. 1C). Next, we assessed C1q protein deposition at the synaptic projection site of the perforant pathway, the hippocampal DGML. In hM3Dq+CNO mice with increased MEC activity, more C1q deposition was observed at the ipsilateral DGML compared to the contralateral DGML (Fig. 1D-E). In contrast, there were no noticeable differences in C1q levels between the ipsilateral and contralateral DGML in control groups (Fig. 1E). The increase of C1q deposition was region-specific to the synaptic termination site of the activated VGLUT1+ perforant pathway in the hM3Dq+CNO mice (fig. S3). C1q levels remained unchanged between hemispheres in non-targeted regions, i.e., hippocampal DG hilus, striatum and cerebellum (fig. S4). These findings suggest that elevated neuronal activity along the perforant pathway increases C1q deposition at synaptic termination sites in the hippocampus.

**Figure 1:**
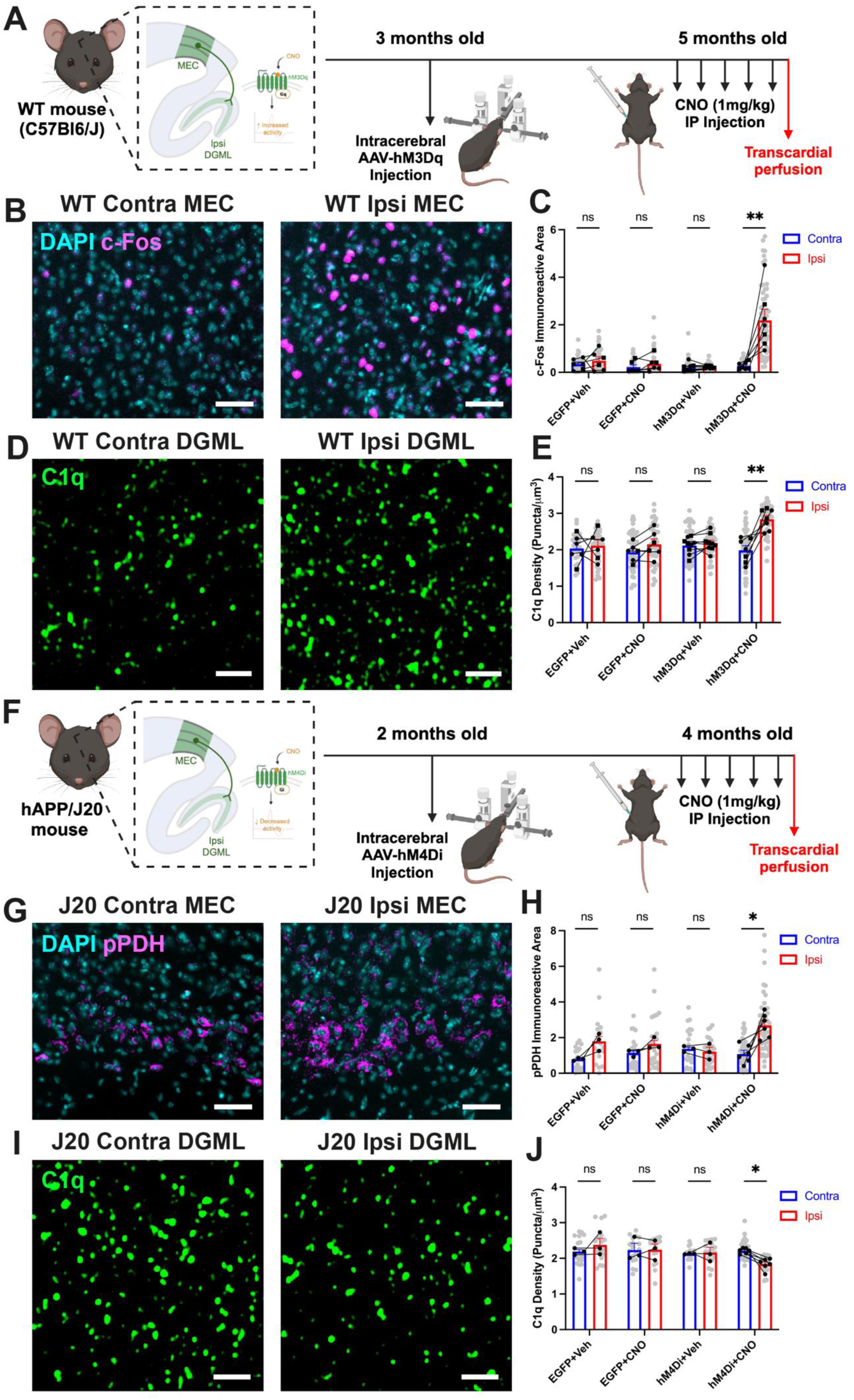
Modulation of neuronal activity alters C1q deposition. **(A)** Schematic of WT experimental paradigm. Created in https://BioRender.com. **(B)** Representative widefield images of c-Fos labeling at contralateral and ipsilateral MEC of hM3Dq+CNO WT mouse, scale bar = 50 μm. **(C)** Quantification of c-Fos immunoreactive area between hemispheres in each mouse group. n = 6 EGFP+Veh mice (contra = 45 ROIs, ipsi = 45 ROIs), n = 5 EGFP+CNO mice (contra = 45 ROIs, ipsi = 45 ROIs), n = 8 hM3Dq+Veh mice (contra = 63 ROIs, ipsi = 63 ROIs) and n = 7 hM3Dq+CNO mice (contra = 66 ROIs, ipsi = 66 ROIs), 2-3 brain sections per hemisphere, 3 ROIs analysed per section. p-values from Bonferroni post-hoc test after 2-way mixed ANOVA with group as between- and hemisphere as within-subject factor (interaction p = 0.008 after log_10_ transformation). **(D)** Representative Airyscan images of C1q puncta at contralateral and ipsilateral DGML of hM3Dq+CNO WT mouse, scale bar = 2 μm. **(E)** Quantification of C1q puncta density between hemispheres in each mouse group. n = 6 EGFP+Veh mice (contra = 33 ROIs, ipsi = 33 ROIs), n = 6 EGFP+CNO mice (contra = 48 ROIs, ipsi = 48 ROIs), n = 9 hM3Dq+Veh mice (contra = 48 ROIs, ipsi = 48 ROIs) and n = 7 hM3Dq+CNO mice (contra = 54 ROIs, ipsi = 54 ROIs), 1-3 brain sections per hemisphere, 3 ROIs analysed per section. p-values from Bonferroni post-hoc test after 2-way mixed ANOVA with group as between- and hemisphere as within-subject factor (interaction p = 0.004). **(F)** Schematic of J20 experimental paradigm. Created in https://BioRender.com. **(G)** Representative widefield images of pPDH labeling at contralateral and ipsilateral MEC of hM4Di+CNO J20 mouse, scale bar = 50 μm. **(H)** Quantification of pPDH immunoreactive area between hemispheres in each mouse group. n = 3 EGFP+Veh mice (contra = 18 ROIs, ipsi = 18 ROIs), n = 3 EGFP+CNO mice (contra = 25 ROIs, ipsi = 26 ROIs), n = 3 hM4Di+Veh mice (contra = 22 ROIs, ipsi = 27 ROIs) and n = 6 hM4Di+CNO mice (contra = 54 ROIs, ipsi = 52 ROIs), 2-3 brain sections per hemisphere, 3 ROIs analysed per section. p-values from Bonferroni post-hoc test after 2-way mixed ANOVA with group as between- and hemisphere as within-subject factor (interaction p = 0.020). **(I)** Representative Airyscan images of C1q puncta at contralateral and ipsilateral DGML of hM4Di+CNO J20 mouse, scale bar = 2 μm. **(J)** Quantification of C1q puncta density between hemispheres in hM4Di+CNO J20 mouse. n = 3 EGFP+Veh mice (contra = 18 ROIs, ipsi = 18 ROIs), n = 3 EGFP+CNO mice (contra = 18 ROIs, ipsi = 18 ROIs), n = 3 hM4Di+Veh mice (contra = 18 ROIs, ipsi = 18 ROIs) and n = 6 hM4Di+CNO mice (contra = 33 ROIs, ipsi = 33 ROIs), 2 brain sections per hemisphere, 3 ROIs analysed per section. p-values from Bonferroni post-hoc test after 2-way mixed ANOVA with group as between- and hemisphere as within-subject factor (interaction p = 0.025). Throughout, square points represent males and circular points represent females, linked points indicate data from the same mouse brain, where points are linked 1 point = 1 hemisphere average = average of brain sections. Data shown as mean ± SEM. ^ns^p > 0.05, *p < 0.05, **p < 0.01, ***p < 0.001.

To further investigate the relationship between neuronal activity and C1q deposition, we performed a complementary experiment to reduce neuronal activity. Using AAV8- CaMKIIa-hM4Di-HA-IRES-mCitrine, we targeted the perforant pathway in J20 mice at 4 months of age, a timepoint of cortical hyperactivity (*7*). J20 mice overexpress hAPP and constitutively secrete hAPP fragments including Aβ (*10*). Mice were treated daily with CNO for five consecutive days (Fig. 1F). In hM4Di-expressing J20 mice, CNO treatment led to elevated levels of phosphorylated pyruvate dehydrogenase (pPDH) in the ipsilateral MEC compared to the contralateral MEC (Fig. 1G-H), a marker indicative of reduced neuronal activity (*13*). Importantly, these mice also exhibited lower C1q deposition in the ipsilateral DGML, the synaptic projection site, compared to the contralateral DGML (Fig 1I-J). In contrast, no differences in pPDH levels in the MEC or in C1q deposition at the DGML were observed in the respective three control groups of mice, i.e., hM4Di-expressing saline-treated mice and AAV8-CaMKIIa-EGFP-expressing mice treated with either CNO or saline (Fig 1H, J). These findings in J20 mice suggest that reducing neuronal activity can partially mitigate hippocampal C1q deposition in this AD mouse model. Together with the experiments using hM3Dq activation in WT mice, these results support a functional link between neuronal activity levels and C1q deposition at synaptic terminals.

#### Changes in neuronal activity influence synaptic density

Next, we asked whether chemogenetic modulation of neuronal activity affects synapse numbers at the termination site. In hM3Dq-activated WT mice, we assessed VGLUT1- and VGLUT2-immunoreactive presynaptic puncta in the DGML, where most perforant pathway terminals contain VGLUT1 (fig. S3A-E) (*14*). No differences in VGLUT1 puncta levels were found between hemispheres in DREADD-activated mice or controls (Fig. 2A-B). However, hM3Dq+CNO-treated WT mice had reduced VGLUT2 density in the ipsilateral DGML compared to the contralateral side (Fig. 2C-D), suggesting that increased neuronal activity leads to refinement of neighbouring VGLUT2+ presynaptic terminals, likely through activity-dependent competition (*15*).

**Figure 2:**
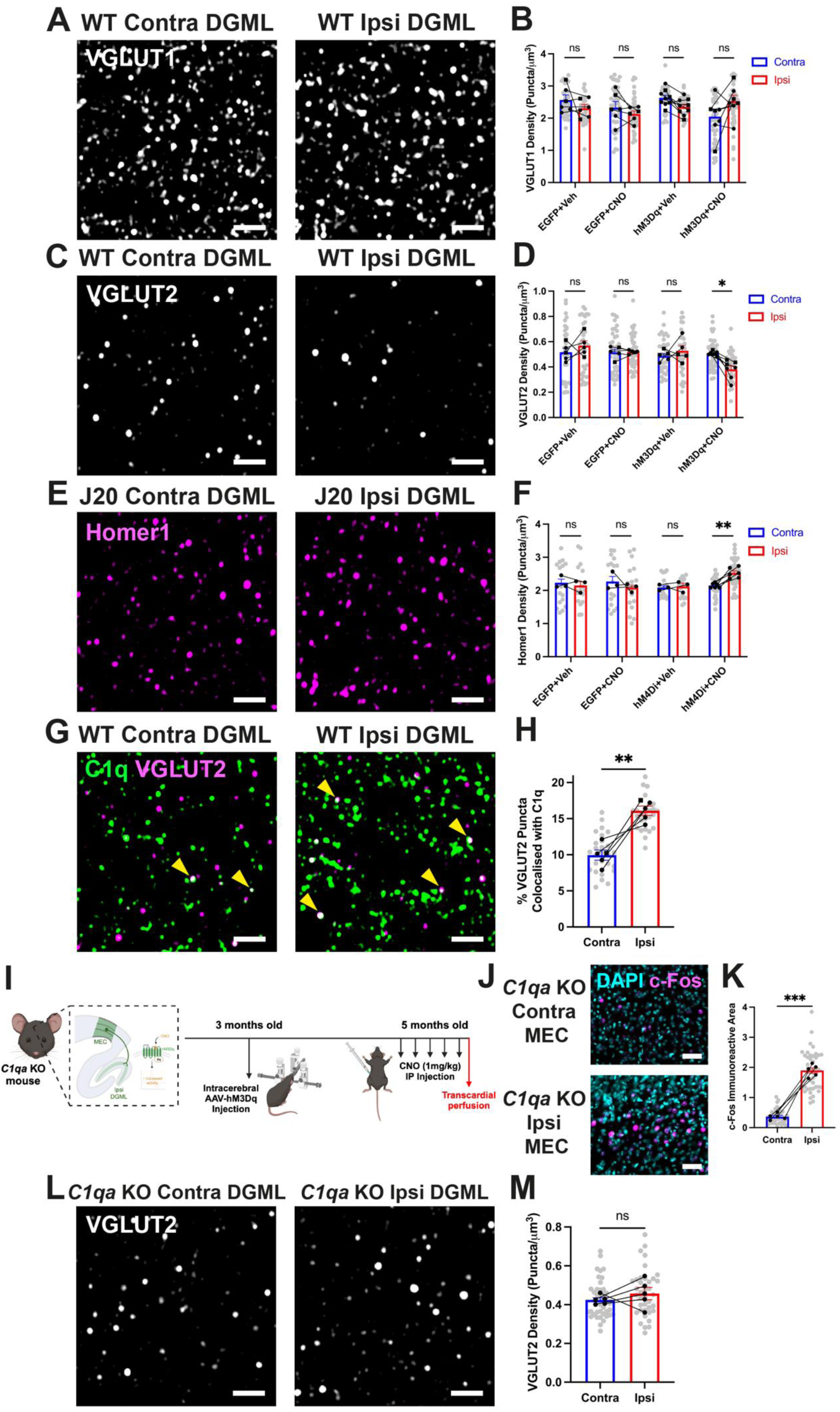
Activity-dependent C1q changes lead to loss of synaptic subsets. **(A)** Representative Airyscan images of VGLUT1 puncta at contralateral and ipsilateral DGML of hM3Dq+CNO WT mouse, scale bar = 2 μm. **(B)** Quantification of VGLUT1 puncta density between hemispheres in each mouse group. n = 6 EGFP+Veh mice (contra = 33 ROIs, ipsi = 33 ROIs), n = 6 EGFP+CNO mice (contra = 48 ROIs, ipsi = 48 ROIs), n = 9 hM3Dq+Veh mice (contra = 36 ROIs, ipsi = 36 ROIs) and n = 7 hM3Dq+CNO mice (contra = 48 ROIs, ipsi = 48 ROIs), 1-3 brain sections per hemisphere, 3 ROIs analysed per section. p-values from Bonferroni post-hoc test after 2-way mixed ANOVA with group as between- and hemisphere as within-subject factor (interaction p = 0.019). **(C)** Representative Airyscan images of VGLUT2 puncta at contralateral and ipsilateral DGML of hM3Dq+CNO WT mouse, scale bar = 2 μm. **(D)** Quantification of VGLUT2 puncta density between hemispheres in each mouse group. Data are mean ± SEM, n = 5 EGFP+Veh mice (contra = 36 ROIs, ipsi = 36 ROIs), n = 5 EGFP+CNO mice (contra = 42 ROIs, ipsi = 42 ROIs), n = 5 hM3Dq+Veh mice (contra = 36 ROIs, ipsi = 36 ROIs) and n = 7 hM3Dq+CNO mice (contra = 39 ROIs, ipsi = 39 ROIs), 2-4 brain sections per hemisphere, 2-3 ROIs analysed per section. p-values from Bonferroni post-hoc test after 2-way mixed ANOVA with group as between- and hemisphere as within-subject factor (interaction p = 0.015). **(E)** Representative Airyscan images of Homer1 puncta at contralateral and ipsilateral DGML of hM4Di+CNO J20 mouse, scale bar = 2 μm. **(F)** Quantification of Homer1 puncta density between hemispheres in hM4Di+CNO J20 mouse. n = 3 EGFP+Veh mice (contra = 17 ROIs, ipsi = 17 ROIs), n = 3 EGFP+CNO mice (contra = 18 ROIs, ipsi = 18 ROIs), n = 3 hM4Di+Veh mice (contra = 15 ROIs, ipsi = 18 ROIs) and n = 6 hM4Di+CNO mice (contra = 40 ROIs, ipsi = 40 ROIs), 2 brain sections per hemisphere, 2-3 ROIs analysed per section. p-values from Bonferroni post-hoc test after 2-way mixed ANOVA with group as between- and hemisphere as within-subject factor (interaction p = 0.014). **(G)** Representative images of C1q and VGLUT2 colocalization at the contralateral and ipsilateral DGML, colocalized puncta indicated by yellow arrowheads, scale bar = 2 μm. **(H)** Quantification of percentage total VGLUT2 puncta colocalizing with C1q at contralateral and ipsilateral DGML of hM3Dq+CNO WT mouse. n = 5 mice (contra = 28 ROIs, ipsi = 27 ROIs), 1 brain section per hemisphere, 3 ROIs analysed per section. p-value from paired t test. **(I)** Schematic of *C1qa* KO experimental paradigm. Created in https://BioRender.com. **(J)** Representative widefield images of c-Fos labeling at contralateral and ipsilateral MEC of hM3Dq+CNO *C1qa* KO mouse, scale bar = 50 μm. **(K)** Quantification of c-Fos immunoreactive area between hemispheres in hM3Dq+CNO *C1qa* KO mouse. n = 5 *C1qa* KO mice (contra = 42 ROIs, ipsi = 42 ROIs), 2-3 brain sections per hemisphere, 3 ROIs analysed per section. p-values from paired t test. **(L)** Representative Airyscan images of VGLUT2 puncta at contralateral and ipsilateral DGML of hM3Dq+CNO *C1qa* KO mouse, scale bar = 2 μm. **(M)** Quantification of VGLUT2 puncta density between hemispheres in hM3Dq+CNO *C1qa* KO mouse. n = 5 *C1qa* KO mice (contra = 39 ROIs, ipsi = 39 ROIs), 2-5 brain sections per hemisphere, 3 ROIs analysed per section. p-values from paired t test. Throughout, square points represent males and circular points represent females, linked points indicate data from the same mouse brain, where points are linked 1 point = 1 hemisphere average = average of brain sections. Data shown as mean ± SEM. ^ns^p > 0.05, *p < 0.05, **p < 0.01, ***p < 0.001.

In 4-month-old J20 mice, (*10*) we observed increased C1q deposition and reduced Homer1-immunoreactive postsynaptic puncta (*8*) in the DGML compared to age- and sex-matched WT mice (fig. S5A-D). Chemogenetic inhibition of the perforant pathway in J20 mice reduced C1q deposition (Fig 1I-J) and increased Homer1 puncta at the ipsilateral DGML (Fig 2E-F). No differences in C1q (Fig 1J) or Homer1 puncta (Fig 2F) were observed between hemispheres in control J20 groups. Together, these results from hM3Dq-activated WT mice and hM4Di-inactivated J20 mice suggest that C1q deposition and loss of synaptic components are, at least partly, activity-dependent.

#### C1q is required for activity-dependent synapse loss in WT mice

In WT mice, activity-dependent VGLUT2 loss was observed in regions corresponding with C1q deposition (Fig 1D). Changes in both C1q and VGLUT2 density were found at the targeted DGML site, but not in the DG hilus, striatum, or cerebellum (fig. S5A-D; fig. S6A-D). The percentage of synapses marked by C1q increases during developmental synapse pruning and in disease states (*8*, *16*). In chemogenetically-activated WT mice, a higher percentage of VGLUT2+ presynaptic terminals colocalized with C1q in the ipsilateral DGML compared to the contralateral side (Fig. 2G-H; fig. S7A), suggesting C1q marks VGLUT2+ synapses for elimination.

To investigate the role of C1q in activity-dependent synapse loss, we performed chemogenetic activation of perforant pathway in *C1qa* KO mice (Fig. 2I) (*17*). Similar to WT mice (Fig 1B-C), hM3Dq-expressing *C1qa* KO mice showed increased c-Fos expression in MEC after CNO treatment (Fig. 2J-K). However, unlike WT mice (Fig. 2C-D), no activity-dependent changes in VGLUT2 puncta in DGML were observed between hemispheres in *C1qa* KO mice (Fig. 2L-M). These results indicate that C1q is required for activity-dependent synapse loss in WT mice.

#### Immunoglobulin-producing B lymphocyte lineage cells are recruited to the site of activity-dependent synapse loss

In the adult brain, C1q is primarily produced by microglia (*18*). As such, single molecule fluorescence *in situ* hybridization (smFISH/RNAScope) showed that *C1qa* mRNA is present in P2Y12+ microglia (Fig S8A). However, despite elevated C1q protein levels in chemogenetically-activated WT mice (Fig 1D-E), *C1qa* mRNA levels did not differ between hemispheres (fig. S8A-B), suggesting that microglia do not increase *C1qa* mRNA production with increased neuronal activity. Additionally, no differences were observed in microglial marker P2Y12, CD68, Iba1, or CD11b immunoreactivity between the DGML of either hemisphere (fig. S8C-H), suggesting no overt microglial inflammation and that other activity-dependent changes may contribute to the increased localization of C1q at hippocampal synapses.

To serve as an unbiased screen for activity-dependent molecular changes at the hippocampal synaptic projection site, we applied spatial transcriptomics (Visium, 10X Genomics) and performed differential gene expression analysis between the ipsilateral DGML and contralateral DGML in chemogenetically-activated WT mice. As expected from the immunostaining data (Fig. S8), Visium data suggested few activity-dependent immune or glial changes at the hippocampal synaptic projection site (Fig 3A). To our surprise, the two gene transcripts upregulated in spots at the ipsilateral DGML of hM3Dq-expressing CNO-treated WT mice were *Igha* and *Igkc* (Fig. 3A-B; fig. S9A-B, S10A-B). Other immunoglobulin gene transcripts were also detectable in the ipsilateral DGML, including *Ighm* and *Jchain* (fig. S10C-D), altogether suggesting that immunoglobulin-secreting B lymphocyte lineage cells localize near the site of activity-dependent synapse loss.

**Figure 3:**
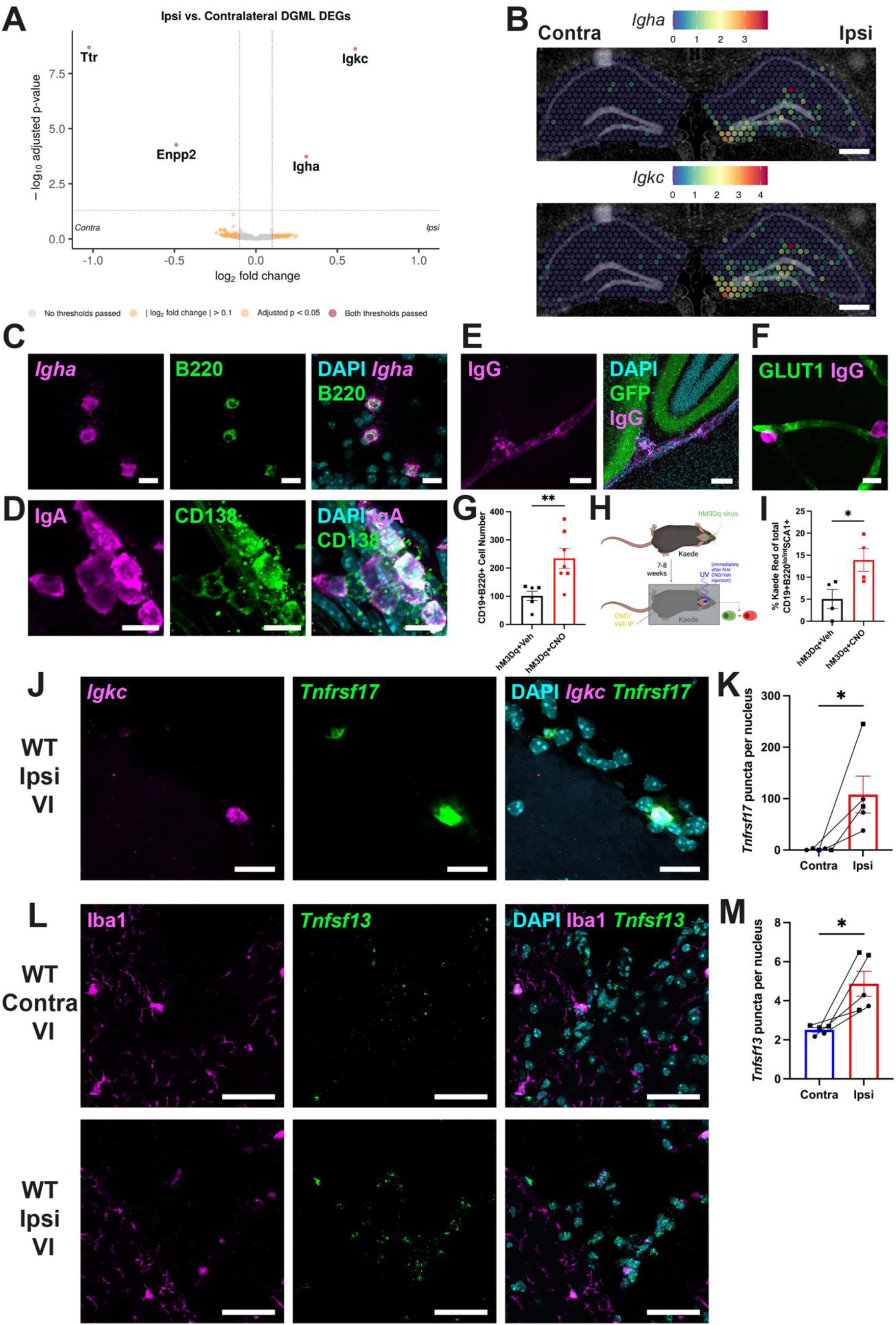
Neuronal activity recruits antigen-experienced B lymphocytes to the hippocampus. **(A)** Volcano plot of DEGs at the DGML spot selection. **(B)** Representative images of Visium spot mRNA transcript counts at contralateral and ipsilateral hippocampi for *Igha* and *Igkc*, scale bar = 500 μm. **(C)** Representative confocal image of *Igha*+ B220+ cells at ipsilateral hippocampal DG, scale bar = 10 μm. **(D)** Representative confocal image of IgA+ CD138+ cells at ipsilateral hippocampal DG, scale bar = 10 μm. **(E)** Representative confocal image of IgG+ cells at ipsilateral hippocampal DG with activated GFP+ projections, scale bar = 100 μm. **(F)** Representative Airyscan image of IgG+ cells juxtaposed to GLUT1+ vessels at ipsilateral hippocampal DG, scale bar = 10 μm. **(G)** Quantification of CD19+B220+ cells at the hippocampus of hM3Dq+Veh and hM3Dq+CNO WT mice by FACS. n = 6 hM3Dq+Veh mice and n = 7 hM3Dq+CNO mice. p-value from unpaired t test. **(H)** Schematic of Kaede experimental paradigm. Created in https://BioRender.com. **(I)** Quantification of percentage Kaede Red+ CD19+B220^lo/int^SCA1+ cells at the hippocampus of hM3Dq+Veh and hM3Dq+CNO Kaede mice by FACS. n = 4 hM3Dq+Veh mice and n = 4 hM3Dq+CNO mice. p-value from unpaired t test. **(J)** Representative confocal images of *Igkc*+ *Tnfrsf17*+ cells at the ipsilateral VI, scale bar = 20 μm. **(K)** Quantification of *Tnfrsf17* puncta per DAPI+ nucleus at the contralateral and ipsilateral VI of the hM3Dq+CNO mouse. n = 5 hM3Dq+CNO mice per group, 1 brain section per hemisphere, 1 ROI analysed per section. p-value from paired t test. **(L)** Representative confocal images of Iba1+ cells and *Tnfsf13* expression at the contralateral and ipsilateral VI of the hM3Dq+CNO mouse, scale bar = 50 μm. **(M)** Quantification of *Tnfsf13* puncta per DAPI+ nucleus at the contralateral and ipsilateral VI of the hM3Dq+CNO mouse. n = 5 hM3Dq+CNO mice per group, 1 brain section per hemisphere, 1 ROI analysed per section. p-value from paired t test. Throughout, square points represent males and circular points represent females, linked points indicate data from the same mouse brain, where points are linked 1 point = 1 hemisphere average = average of brain sections. Data shown as mean ± SEM. *p < 0.05, **p < 0.01.

Using smFISH/RNAScope and immunohistochemistry, we validated the presence of *Igha* and *Igkc*-expressing cells (Fig. 3C; fig. S10E-F), as well as IgA+B220+ B cell cells (Fig. 3C) and IgA+CD138+ plasma cells (Fig. 3D). In particular, IgA+, IgM+ and IgG+ cells were observed where the hippocampal DG intersects with the ventricular space, nearby the hippocampal fissure and at the leptomeningeal velum interpositum (VI) adjacent to the activated DG (Fig. 3D, E; fig. S10G-H). Using Airyscan confocal imaging and 3D reconstruction, we observed that some IgG+ cells exhibited perivascular localization, juxtaposed to GLUT1+ hippocampal blood vessels (Fig. 3F). The region-specific presence of CD19+B220+ B cells was further confirmed using flow cytometry, with a higher B cell count detected in mice bilaterally expressing hM3Dq at the MEC treated with CNO compared to those treated with vehicle (Fig. 3G fig. S11A). In contrast, we did not observe changes in the number of CD19+B220+ B cells in the non-hippocampal brain tissue nor in dural meninges between hM3Dq+CNO and hM3Dq+Veh mouse groups (fig. S11B-C). Intracardiac injection of an anti-CD45 antibody in WT hM3Dq+CNO mice three minutes before perfusion to label circulating immune cells, did not stain hippocampal B cells (fig. S11D), suggesting that these cells are extravascular and located within the tissue. Altogether, these data suggest that B lymphocyte lineage cells localise to the leptomeningeal and perivascular areas adjacent to the site of activity modulation.

To assess whether hippocampal B lymphocyte lineage cells originate from the dural meninges or skull bone marrow (*19*, *20*), we used Kaede transgenic mice, which allow the trafficking of photoconverted cells following UV light exposure (Fig. 3H) (*21*). Upon UV illumination of the skull, we detected a greater proportion of photoconverted CD19+B220^lo^SCA1+ plasmablasts in the hippocampus of the hM3Dq+CNO-injected Kaede mice compared to non-activated controls (Fig. 3I; fig. S11E). In contrast, while photoconversion occurred, there were no differences in the proportions of photoconverted plasmablasts in the dural meninges or non-hippocampal brain regions (fig. S11F-G). These results suggest that a proportion of the B lymphocyte lineage cells found at the hippocampus following chemogenetic activation originate from the skull bone marrow and/or dural meninges.

Plasma cells require niche cytokines, such as A proliferation-inducing ligand (APRIL; gene name *Tnfsf13*), to sustain their survival in tissues (*22–25*). We observed increased expression of *Tnfsf13* at the ipsilateral compared to the contralateral, non-activated VI (Fig. 3L-M). Furthermore, *Tnfrsf17, encoding* B cell maturation antigen (BCMA), a receptor expressed by mature B cells and plasma cells, was also detectable in *Igkc*-expressing cells at the ipsilateral VI, nearby the activated DG (Fig. 3J-K). Overall, these findings suggest that mature B lymphocyte lineage cells localize to the hippocampus in an activity-dependent manner.

#### Immunoglobulins may be required for activity-dependent C1q deposition and synapse loss in the adult WT brain

We next sought to determine the functional relevance of B lymphocyte lineage cells in the activated hippocampus. B cell receptor sequencing (BCRseq) of dural meningeal, hippocampal and striatal (negative control) tissues from hM3Dq+CNO mice showed a greater number of BCRs in hippocampal tissue compared with striatum (Fig. 4A-B), in keeping with the spatial transcriptomic and confocal imaging data. Several B cell clones included BCRs from all three tissues, with a higher number of shared BCRs between the dural meninges and the hippocampus than the dura and striatum (Fig. 4A-B), supporting a meningeal origin for the hippocampal B lymphocyte lineage cells, consistent with the Kaede photoconversion study. The BCRs present in the hippocampus of hM3Dq+CNO mice included IgA, IgG, and IgM heavy chains (Fig. 4C), in keeping with our confocal imaging data.

**Figure 4:**
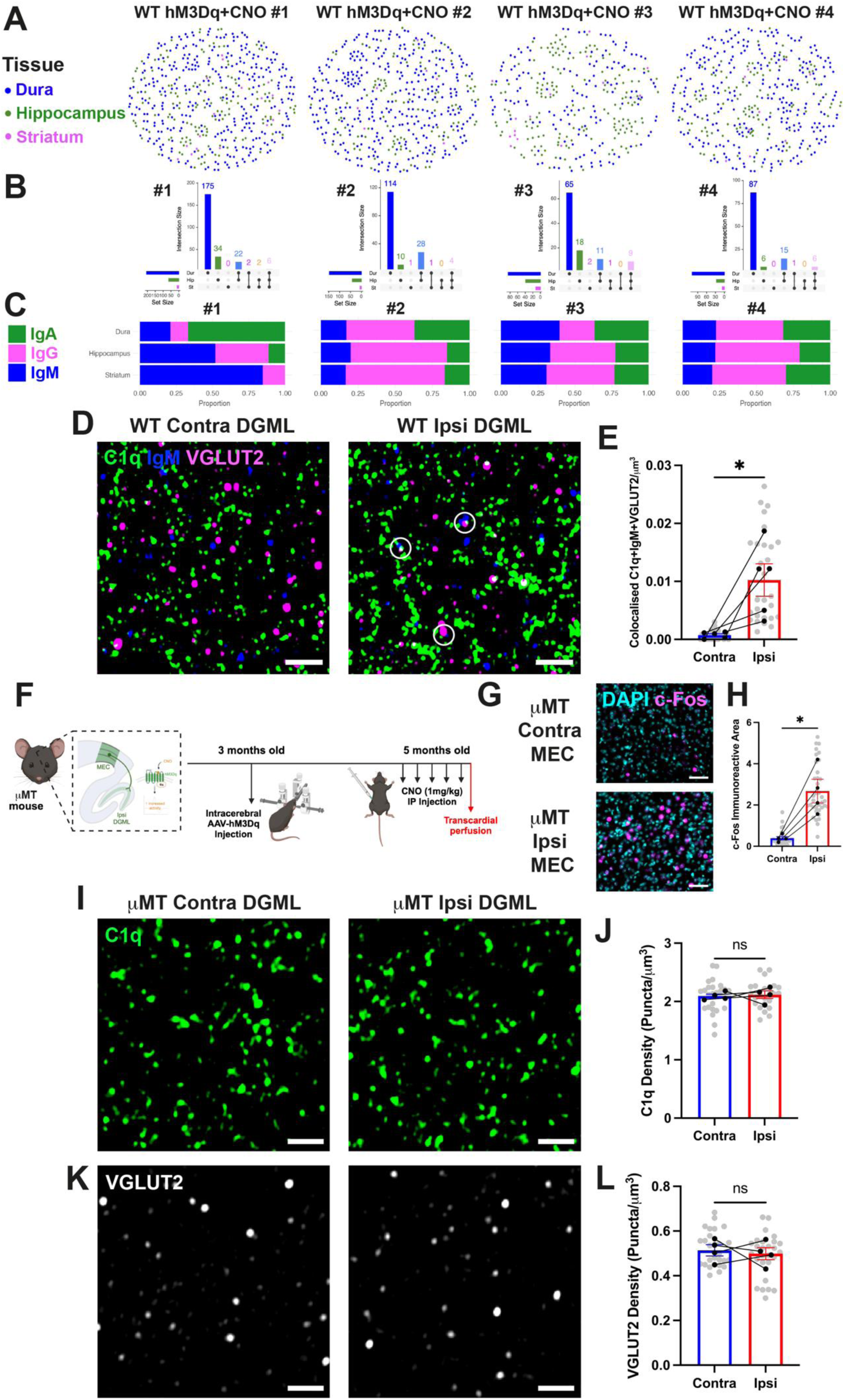
B lymphocyte lineage cells produce IgM to facilitate activity-dependent synapse loss. **(A)** BCR network plots for separate hM3Dq+CNO mice coloured by tissue distribution. Each circle/node corresponds to a single BCR contig and connected circles/nodes indicate that contigs are similar. **(B)** Upset plots of BCR contig number. The vertical graphs show the number of unique and overlapping BCR contigs for each tissue with the intersection indicated by the dots connected by lines below the X-axis. The number of BCR contigs is indicated above the columns. The horizontal graphs on the left show the total number of BCR contigs detected in each tissue respectively. **(C)** Bar graphs demonstrating the proportion of each heavy chain isotype class in the total BCR repertoire for different tissues. **(D)** Representative Airyscan images of IgM puncta colocalized with double positive C1q+VGLUT2+ puncta at contralateral and ipsilateral DGML of hM3Dq+CNO WT mouse, scale bar = 2 μm. **(E)** Quantification of colocalized IgM, C1q and VGLUT2 puncta between hemispheres in hM3Dq+CNO WT mouse. 5 hM3Dq+CNO mice (contra = 28 ROIs, ipsi = 28 ROIs), 2 brain sections per hemisphere, 2-3 ROIs analysed per section. p-values from paired t test. **(F)** Schematic of μMT experimental paradigm. Created in https://BioRender.com. **(G)** Representative widefield images of c-Fos labeling at contralateral and ipsilateral MEC of hM3Dq+CNO μMT mouse, scale bar = 50 μm. **(H)** Quantification of c-Fos immunoreactive area between hemispheres in hM3Dq+CNO μMT mouse. n = 4 μMT mice (contra = 36 ROIs, ipsi = 36 ROIs), 3 brain sections per hemisphere, 3 ROIs analysed per section. p-values from paired t test. **(I)** Representative Airyscan images of C1q puncta at contralateral and ipsilateral DGML of hM3Dq+CNO μMT mouse, scale bar = 2 μm. **(J)** Quantification of C1q puncta density between hemispheres in hM3Dq+CNO μMT mouse. 4 μMT mice (contra = 24 ROIs, ipsi = 24 ROIs), 2 brain sections per hemisphere, 3 ROIs analysed per section. p-values from paired t test. **(K)** Representative Airyscan images of VGLUT2 puncta at contralateral and ipsilateral DGML of hM3Dq+CNO μMT mouse, scale bar = 2 μm. **(L)** Quantification of VGLUT2 puncta density between hemispheres in hM3Dq+CNO μMT mouse. 4 μMT mice (contra = 27 ROIs, ipsi = 27 ROIs), 2-3 brain sections per hemisphere, 3 ROIs analysed per section. p-values from paired t test. Throughout, circular points represent females, linked points indicate data from the same mouse brain, where points are linked 1 point = 1 hemisphere average = average of brain sections. Data shown as mean ± SEM. ^ns^p > 0.05, *p < 0.05.

While it is well-established that classical complement cascade activation involves C1q fixation to IgG or IgM (*26*, *27*) outside of the CNS, the role of immunoglobulins within the CNS has largely been overlooked due to their limited presence in homeostasis. Our identification of B/plasma cells at sites of increased C1q deposition prompted us to investigate whether immunoglobulins are present on C1q+ VGLUT2 presynaptic terminals in the chemogenetically-activated brain. Using Airyscan super-resolution confocal microscopy, we observed an increase in IgM-immunoreactive puncta on C1q+VGLUT2+ terminals at the activated projection site (ipsilateral DGML) compared to the non-activated site (contralateral DGML) (Fig. 4D-E). In contrast, we did not detect punctate staining for IgA or IgG (data not shown). These results suggest that IgM may facilitate C1q binding to VGLUT2+ presynaptic terminals, akin to its role outside the brain (*28*).

To address whether the presence of B/plasma cells is necessary for activity-dependent C1q deposition and VGLUT2 loss, we expressed hM3Dq in perforant pathway projection neurons of μMT mice, which lack the IgM heavy chain and are therefore B and plasma cell-deficient, and lack secreted immunoglobulins (Fig. 4F) (*29*). As observed in WT mice, hM3Dq+CNO treatment led to an increase in c-Fos expression in the ipsilateral MEC of μMT mice (Fig. 4G-H), indicating activation of the perforant pathway. However, there were no activity-dependent changes in C1q deposition (Fig 4I-J) or VGLUT2 presynaptic puncta density (Fig. 4K-L) between the ipsilateral and contralateral DGML in chemogenetically-activated μMT mice. Altogether, these results suggest that IgM is required for activity-dependent C1q deposition and synapse loss in hM3Dq-activated WT mice.

To further explore the role of B lymphocyte lineage cells and immunoglobulin in activity-dependent C1q deposition, we used kainic acid (KA) at a subthreshold dose (14mg/kg, administered intraperitoneally) that avoids generalized seizures or cell death (*30*). As expected, KA injection increased hippocampal c-Fos expression (fig. S12A-B), confirming elevated neuronal activity. In WT mice, daily KA injections over three consecutive days led to increased C1q deposition in the hippocampal CA3 stratum lucidum (CA3SL; fig. S12C-D), similar to the effects observed in the hM3Dq chemogenetic model (Fig 1D-E). In contrast, KA did not induce C1q deposition in μMT mice (fig. S12E-F), further supporting the conclusion that B/plasma cells and/or their secreted immunoglobulins are required for KA-induced C1q deposition in the context of neuronal hyperactivity.

## Discussion

Neuronal activity regulates synapse pruning during development (*31*) and microglia-mediated synapse elimination (*1*), but its role in region-specific C1q deposition and synapse loss in the adult brain, beyond developmental synaptic pruning, has been less clear. Using *in vivo* chemogenetics, we show that heightened cortical activity along the perforant pathway leads to C1q deposition and synapse loss in the hippocampus. These findings identify neuronal hyperactivity as a mediator of C1q deposition at synapses in the adult brain. Interestingly, we observe a selective elimination of VGLUT2+ presynaptic terminals while the activated VGLUT1+ neuronal projections are spared, a pattern that echoes competition-dependent synapse refinement during development (*15*). Future studies are warranted to elucidate precise molecular mediators underlying activity-dependent synapse loss in the adult brain. Activity-dependent cues, such as “eat-me” and “don’t-eat-me” signals on synapses (e.g., PirB (*32*), CD47 (*33*), phosphatidylserine (*34*)), may play a role in this process. Alternatively, microglia and astrocytes which provide critical feedback mechanisms in response to neuronal hyperexcitability (*35–37*), may regulate this process. Although we controlled for various factors such as surgery-induced inflammation, cell death and generalized seizures, neuronal hyperactivity could induce low levels of excitotoxicity (*38*), leading to generation of local damage signals at sites of synaptic refinement.

Our study suggests a functional link between neuronal hyperactivity and C1q- mediated synapse loss in the adult CNS, raising the possibility that dysregulated neuronal activity, an early hallmark of AD (*39*, *40*), contributes to region-specific C1q deposition and synapse loss (*8*). In early stages of AD, a pathological feedback loop likely forms between neuronal hyperactivity and synaptotoxic β-amyloid (Aβ), where neuronal activity promotes Aβ and/or tau production at the synapse (*41–44*), leading to complement deposition and increased synapse vulnerability; in turn, Aβ aggregates further exacerbate dysregulated neuronal activity (*40*, *45*). Our findings indicate that reducing neuronal hyperactivity along the perforant pathway in 4-month-old J20 mice alleviates hippocampal C1q deposition and partially restores Homer1-immunoreactive synaptic puncta. This beneficial effect may stem from decreased levels of Aβ and/or tau, as reduced neuronal activity in the perforant pathway has been associated with lower hippocampal Aβ and tau levels (*46*, *47*). Consistent with this, the antiepileptic drug levetiracetam alleviates synaptic and cognitive deficits in J20 and APP/PS1 mouse models of AD (*48*, *49*) and reduces hippocampal hyperactivation while improving hippocampal-dependent memory function in individuals with amnestic mild cognitive impairment (*50*). These studies suggest that neuronal hyperactivity may be an early determinant of synapse vulnerability, contributing to degeneration and loss. Thus, strategies to suppress neuronal hyperactivity may prove beneficial, particularly in subgroups of AD patients exhibiting epileptic- or seizure-like symptoms (*51*).

Unexpectedly, unbiased spatial transcriptomics implicated local immunoglobulin production in activity-dependent C1q deposition. Using live-cell tracking, BCRseq, super-resolution microscopy, and *in vivo* chemogenetics in genetic KO mice, we show that neuronal hyperactivity leads to accumulation of B cells and plasma cells in the hippocampus, a site of activity-dependent synapse loss in our model. These B lymphocyte lineage cells, which are present in meningeal immune niches (*19*, *20*), appear to migrate to the VI near the hippocampus upon hyperactivity, where their presence may be stabilized via APRIL-BCMA signaling, and may contribute to IgM- mediated complement deposition. One caveat of the μMT model used for functional analysis is that these mice are not only IgM-deficient, but also lack mature B and plasma cells, meaning that antibody-independent B cell functions such as pro- and anti-inflammatory cytokine production (*52*, *53*) are also lacking in these mice. While IgM is well-established in non-CNS tissues to bind and activate C1q for immune clearance (*28*), the role of immunoglobulins and their interaction with C1q in the CNS has been underexplored. Here, we show the functional relevance of B lymphocyte lineage cells using two distinct paradigms of neuronal hyperactivity, suggesting a previously unrecognized role for humoral immunity in activating C1q for synapse elimination within the CNS.

The adaptive immune system’s role in the CNS, particularly its interactions with microglia and the innate immune system, remains largely unexplored. Here we propose a mechanism by which intercellular crosstalk between C1q and immunoglobulins facilitates synapse loss upon neuronal hyperactivity. The relevance of immunologublin-C1q interaction in disease remains to be determined. Interestingly, a proteomic study suggested upregulation of IgG3 and C1q on synapses of AD postmortem brain tissue (*54*). Furthermore, in models of Guillain-Barré syndrome, anti-ganglioside IgM and IgG3 antibodies interact with C1q at neuromuscular junction axon terminals to facilitate peripheral neuropathy (*55*), suggesting that immunoglobulin-mediated antigen recognition can activate C1q on synaptic terminals. These findings suggest the potential presence of immunoglobulin-C1q complexes on synapses in diseased brain. However, further research in relevant disease models is needed to confirm this and explore potential links to neuronal activity.

Future studies are required to elucidate the underlying mechanisms and relevance of activity-induced B and/or plasma cell recruitment. Key questions remain: What antigens are the immunoglobulins recognizing? Does B and/or plasma cell recruitment occur during normal physiology or only when aberrant neuronal hyperactivity occurs? Do similar innate-adaptive interactions extend to circuits in different brain regions which may not be as overtly plastic or as easily accessible to immune niches as the hippocampus? (*56*) Furthermore, natural antibodies, which are germline-encoded immunoglobulins targeting self-antigens, bind phosphatidylserine and activate C1q to facilitate removal of apoptotic bodies (*57*). Similar apoptotic signals including phosphatidylserine are exposed on synapses undergoing elimination (*34*, *58*), raising questions about the presence of natural antibodies in the CNS and their involvement in activity-driven synapse loss. Future work will seek to clarify their role and whether their presence in CNS immune niches is conserved.

Altogether, our findings demonstrate that neuronal hyperactivity serves as a trigger for C1q deposition at synaptic termination sites and subsequent synapse loss in the adult brain, with immunoglobulin deposition potentially contributing to this process. Specifically, we propose innate-adaptive immune crosstalk facilitating synapse loss, suggesting previously unexplored interplay between neuronal activity, immune signaling, and synaptic fate in the CNS. Given the relevance of neuronal hyperactivity in neurodegenerative diseases (*59*) and C1q in synaptopathy (*60*), our study may have broad implications for synaptic dysfunction in neurologic diseases.

## Acknowledgements

We thank members of the Hong lab and members of the International Neuroimmune Consortium for critical input to the manuscript; Elena Ghirardello and Phillip Muckett for managing mouse colonies; Carlo Sala Frigerio for guidance with Visium methodology; Nicholas Cade for microscopy; Ryan Wee (UCL) for stereotaxic surgery training; Gustavo A. Rodriguez, S. Abid Hussaini (Columbia University, New York) and Karen E. Duff (UKDRI) for providing tissue and guidance on viral targeting of MEC; Leen Ali and Kiavash Movahedi (VIB Center for Inflammation Research, Brussels) for preliminary IgG/CD138 immunostaining; Jorge J. Palop and Keran Ma (Gladstone Institute, San Francisco) for providing tissue for preliminary C1q immunostaining; Alec Walker (Boston Children’s Hospital and Harvard Medical School) for guidance on B lymphocytes, FACS and Kaede experiments; Jasmin Herz and Jonathan Kipnis (Washington University, St Louis) for guidance on Kaede experiments and Sandro Da Mesquita (Mayo Clinic, Jacksonville) for guidance on B lymphocytes at brain borders.

## Funding

UK DRI Ltd, funded by the UK Medical Research Council, Alzheimer’s Society and Alzheimer’s Research UK grant UKDRI-1011 and UKDRI-1209 (SH)

Chan Zuckerberg Collaborative Pairs Initiative DAF2022-250425 (SH)

Anonymous Foundation (SH)

Alzheimer’s Association Research Grant 23AARG-1018881 (SH)

Wellcome Trust studentship (219906/Z/19/Z) (GC)

Wellcome Trust Sir Henry Wellcome Fellowship (221634/Z/20/Z) (SDS)

Wellcome Trust and the Royal Society 109360/Z/15/Z (AFM)

Medical Research Council MR/W02005X/1 (AFM)

Wellcome Trust Investigator Award 220268/Z/20/Z (BT, TS, MRC)

Wellcome Discovery Award 227890/Z/23/Z (MRC)

## Author contributions

Conceptualization: GC, SH

Methodology: GC, ET, SDS, BJYT, DP, AM, MC, SH

Investigation: GC, ET, MK, SDS, BJYT, JRC, MT, JCF, JZG, GY, CP, DP, LSSF, DS, VK

Validation: ET, MK, SDS, MT, JCF, JZG, GY

Formal analysis: GC, ET, MK, SDS, BJYT, MT, JCF, JZG, GY, AS

Resources: CSF, AFM, MRC, SH

Data Curation: ET

Funding Acquisition: GC, SH

Visualization: GC, SH

Project administration: MRC, SH

Supervision: GC, MRC, SH

Writing - Original draft: GC, MRC, SH

## List of Supplementary Materials

Materials and Methods

Figs. S1 to S12

Table S1 to S3

References (*61–76*)

## Materials and Methods

### Animals

All animal experiments were performed in accordance with the UK Animals (Scientific Procedures) Act (ASPA) 1986. Experimental procedures were approved by the UK Home Office (Home Office license 70/8999) and following local ethical advice by consultation with University College London (UCL) veterinary staff. All animal work was carried out under the appropriate ethics and licences.

Mice were housed in individually ventilated cages on a 12-hour light/dark cycle in a temperature- and humidity-controlled environment (23.1 °C, 30-60% humidity) under pathogen-free conditions with ad libitum access to water and food at all times. C57BL/6J wild-type (WT) mice were obtained from Charles River UK. All mouse strains were bred on a C57BL/6J background. hAPP-J20 transgenic mice (J20; B6.Cg- Zbtb20Tg (PDGFB-APPSwInd)20Lms/2Mmjaxn) (*10*) were kindly provided by Lennart Mucke, University of California, San Francisco and distributed by Patricia Salinas, UCL. *C1qa*-/- mice (*C1qa* KO) (*17*) were kindly provided by Marina Botto, Imperial College London. *Ighm*-/- mice (μMT or *Ighm* KO) (*29*) were obtained from JAX (C57BL/6.129S2-Ighmtm1Cgn/J, strain #002288) via the Francis Crick Institute central mouse repository.

For all experiments, appropriate sex- and age-matched controls were used. Both male and female mice were used throughout the experiments presented and details of mouse sex are included in graphs and figure legends. Pilot experiments were conducted using both female and male mice to ensure there were no sex-specific phenotypes. Mice were housed individually for at least one month after surgical procedures to avoid compromising sutures. Genotyping for all mouse lines was performed by Transnetyx using tissue from ear punch samples taken to identify mice. For all experiments, the appropriate genotype-, sex- and age-matched controls were used.

For transcardial perfusion and tissue harvest, mice were deeply anaesthetized by IP injection of a sodium pentobarbital (200 μg/g body weight; Dolethal, Vetoquinol) overdose. Upon loss of hindlimb and tail pinch reflexes, the abdomen was opened and the heart exposed. A needle was inserted into the left ventricle of the heart and the right atrium punctured using scissors. 20 mL ice-cold filtered phosphate-buffered saline (PBS; 18912, Thermo Fisher Scientific) was pumped at a steady rate through the circulation followed by 20 mL ice-cold ultrapure 4% paraformaldehyde (PFA; 18814-20, Generon; diluted in 0.2 μm-filtered PBS). After perfusion, the mouse head was removed and the brain carefully extracted from the skull followed by post-fixation for 24 hours in 4% PFA at 4 °C.

### Stereotaxic surgery

Stereotaxic surgeries were performed as previously described (*61*, *62*). Briefly, adult mice (3mo for WT hM3Dq experiments, 2mo for J20 hM4Di experiments) were anaesthetized with isoflurane (IsoFlo, Zoetis) at 4% in oxygen for induction. Mice were head-fixed into a digital stereotaxic frame (504926, World Precision Instruments) and anaesthesia was maintained at 1.5-2% isoflurane concentration in 1.5 L/min oxygen flow. Aseptic conditions were adopted throughout. Sterile liquid gel drops (Viscotears, Bausch and Lomb) were applied to keep the eyes of anaesthetized mice moist. After cleaning the scalp with a 1:1 solution of 0.9% saline (Aqupharm 3, Animalcare):chlorhexidine gluconate (4% w/v Hibiscrub, Regent Medical), bupivacaine local anaesthesia (0.025% Marcain Polyamp, AstraZeneca) was applied and a single skin incision along the scalp midline exposed the skull. Bregma was identified and the skull levelled in the mediolateral (ML) and anteroposterior (AP) aspects. Sterile 0.9% saline was applied at regular intervals to prevent the skull and connective tissue from excessive drying. A craniotomy was drilled using an OmniDrill35 Micro Drill (503599, World Precision Instruments) with 0.9 mm burr (#330104001001007, Hager and Meisinger) unilaterally (always right hemisphere) at stereotaxic coordinates AP −4.60, ML 3.00, dorsoventral (DV) −3.00 (all in mm; from Paxinos and Franklin’s The Mouse Brain in Stereotaxic Coordinates, Fourth Edition) relative to bregma to target the MEC. Pulled long-shaft borosilicate pipettes (3.5” tubes, #3-000-203-G/X, Drummond Scientific) were backfilled with mineral oil (330779, Sigma-Aldrich) before being loaded with either DREADD virus (AAV8- CaMKIIa-HA-hM3Dq-IRES-mCitrine, #50466-AAV8, Addgene; titre = 1.6×10^13^ gc/mL or AAV8-CaMKIIa-HA-hM4Di-IRES-mCitrine, #50467-AAV8, Addgene; titre = 1.6×10^13^ gc/mL) or control EGFP virus (AAV8-CaMKIIa-EGFP, #50469-AAV8, Addgene; titre = 2.3×10^13^ gc/mL). Controlled injection of 200 nL virus at 80 nL/min rate was conducted using a NanoFil 10 μL syringe (World Precision Instruments) connected to an UltraMicroPump-3 (World Precision Instruments) via NanoFil injection holder (World Precision Instruments) and SilFlex tubing (World Precision Instruments). The pipette was left in place for 10 minutes post-injection followed by slow retraction to prevent the backflow of contents. The scalp incision was sutured using coated Vicryl polygalactin 910 undyed braided absorbable sutures with 11 mm 3/8c reverse cutting Ethalloy Prime hook (W9500, Ethicon) and closed with Vetbond tissue adhesive (1469SB, 3M) to seal the scalp incision. After incision closure, mice were allowed to recover post-surgery for one hour in a heated chamber. Subcutaneous carprofen (5 μg/g body weight; Carprieve, Norbrook) and buprenorphine (0.1 μg/g body weight; Vetergesic, Ceva) diluted in sterile 0.9% saline were administered peri-operatively and carprofen was added to the drinking water (33.33 mg/mL) for 3 days post-surgery.

### Substance administration

Kainic acid (KA; 14 mg/kg body weight; K0250, Sigma-Aldrich) dissolved in 0.9% saline or equal volume 0.9% saline vehicle were administered IP daily on three consecutive days followed by transcardial perfusion 90 minutes after the final injection. For 90 minutes after KA injections, mice were monitored for seizure-like activity and scored on a modified Racine scale (*63*, *64*) (see Table 2.1). For DREADD experiments, IP CNO administration began a minimum of seven weeks after AAV delivery. A 5 mg/mL working solution of CNO dihydrochloride (water-soluble; HB6149, Hello Bio) dissolved in 0.9% saline was stored at −80 °C until aliquoting and 100 μL aliquots were stored at −20 °C until day of use. Sufficient volume of 5 mg/mL CNO solution was diluted in 0.9% saline after weighing mice just before the first IP dose to generate a concentration allowing a dose of 1 mg CNO/kg body weight with a 100 μL volume injection. 100 μL diluted CNO solution or 100 μL 0.9% saline vehicle were administered IP daily on five consecutive days. Mice were transcardially perfused 90 minutes after the final CNO or vehicle injection. Mice administered CNO were not observed to display seizure-like activity as measured on the modified Racine scale.

### Immunohistochemistry (IHC)

Brains were washed thoroughly in PBS for at least two hours at room temperature (RT) followed by incubation for at least two days in 30% sucrose (S0389, Sigma-Aldrich) solution at 4 °C. After sinking in sucrose solution, brains were embedded in optimal cutting temperature (OCT) compound (4583, Tissue-Tek) and frozen in moulds using 2,3-methylbutane (M32631, Sigma-Aldrich) on dry ice. Horizontal and sagittal sections were cut on a cryostat. 15 μm-thick sagittal sections (0.3-0.6μm in mediolateral plane) were placed directly onto Superfrost Plus GOLD glass slides (K5800AMNZ72, Thermo Fisher Scientific) for some IHC and all RNAscope experiments while 30 μm-thick free-floating sagittal (0.6-1.5μm in mediolateral plane) and horizontal sections (2.1-3.3μm in dorsoventral plane) were placed in wells of 24-well plates with 1.5 mL cryoprotectant solution (40% PBS, 30% ethylene glycol [324558, Sigma-Aldrich], 30% glycerol [G7757, Sigma-Aldrich]) using stainless steel forceps. 10 sections were stored in each well, meaning each well spanned 0.3 mm in each plane of sectioning, ML for sagittal sections and DV for horizontal sections. Sections on glass slides were stored at −80 °C and free-floating sections were stored in cryoprotectant at −20 °C until IHC.

Free-floating sections were left shaking on an orbital shaker at ∼90 rpm during all washing, blocking and antibody incubation steps. Washing consisted of a minimum of three 10-minute washing steps in PBS. If a primary mouse antibody was used, M.O.M.® (Mouse on Mouse) Blocking Reagent (Vectorlabs, MKB-2213-1) was added before blocking. Sections for immunostaining were first washed, then blocked and permeabilized for two hours at room temperature (RT) in 20% normal goat serum (NGS; ab7481, Abcam) (20% normal donkey serum [NDS; ab7475, Abcam] used instead when IgA included) and 1% Triton X-100 (28314, Thermo Fisher Scientific) diluted in PBS. Primary antibodies were incubated overnight at 4 °C in 10% NGS and 0.5% Triton X-100 in PBS. Washing was followed by secondary antibody incubation in PBS with 10% NGS and 0.5% Triton X-100 for two hours at RT. Sections were again washed before 1:10000 DAPI (D1306, Thermo Fisher Scientific) staining for 10 minutes prior to mounting.

Sections for synapse marker (Bassoon+Homer1) immunostaining were first washed then pre-treated with 1% Triton X-100 before blocking. These sections were then blocked and permeabilized for two hours at RT in 20% NGS, 1% bovine serum albumin (BSA; A2153, Sigma-Aldrich) and 0.3% Triton X-100 in PBS. Primary antibodies were incubated overnight at 4 °C in 20% NGS, 1% BSA and 0.3% Triton X-100 in PBS. Wash steps preceded secondary antibody application at 1:500 in 20% NGS, 1% BSA and 0.3% Triton X-100 for four hours at RT. Washing was followed by 1:10000 DAPI staining for 10 minutes before a final wash and mounting.

For c-Fos immunostaining, sections were washed then blocked for one hour at RT in 10% horse serum (26050-070, Thermo Fisher Scientific) and 1% Triton X-100 in PBS. Primary and secondary antibodies were also incubated in 10% horse serum and 1% Triton X-100 in PBS at 4 °C overnight and at RT for two hours respectively. Washing steps were as in previous protocols and 1:10000 DAPI was applied for 10 minutes before mounting. All slides were mounted on Superfrost glass slides (ISO 8037/1, Epredia) using Prolong Gold Antifade mountant (P36930, Thermo Fisher Scientific) with 22 x 50 mm #1.5 coverslips (15797582, Menzel-Glaser).

### *In situ* hybridization (RNAscope)

RNAscope *in situ* hybridization (ISH) experiments were performed as per the manufacturer’s specifications using the RNAscope Fluorescent Multiplex Assay (320293, Advanced Cell Diagnostics Bio) in an RNase-free environment. Briefly, 15 μm brain sections on Superfrost Plus GOLD slides (Thermo Fisher Scientific, K5800AMNZ72) were first incubated in hydrogen peroxide (1.07209, Sigma-Aldrich) for four minutes at RT and then washed in RNase-free water. Slides underwent target antigen retrieval at boiling point for four minutes and dehydration in 100% ethanol for five minutes followed by Protease Plus treatment for 15 minutes at RT before probes against the target gene (*C1qa*: 441221-C2, *Igha*: 414281-C2, *Igkc*: 414291-C2, *Tnfsf13*: 547331-C1, *Tnfrsf17*: 414871-C1), all Advanced Cell Diagnostics Bio) hybridization for two hours at 40 °C. Alternating amplification and wash steps followed before addition of Opal 570 fluorophore. Slides were then processed using the IHC protocol above for staining of P2Y12 microglia marker or B220 B cell marker and mounting of slides.

For *C1qa* mRNA analysis, maximum intensity projections were generated on ImageJ before conversion to .ims files. For *C1qa* counting within P2Y12+ cells, number of *C1qa* puncta within P2Y12 channel area of maximum intensity projections was calculated.

### FACS Experiments

To obtain single cell suspensions from hippocampus, brains were first quickly isolated from the skull and placed on ice. The dorsal part of the skull was carefully removed, meningeal dura mater was peeled off the skull cap and hippocampus was dissected from the rest of the brain tissue on ice using chilled instruments. The tissues were finely chopped using chilled razorblades and transferred to tubes containing ice-cold RPMI-1640 supplemented with 10% FBS and 20mM HEPES medium. Single-cell suspensions from brain tissue were prepared using the Dissociation kit (Miltenyi Biotech) according to the manufacturer’s instructions. Tissue chunks were pelleted by centrifugation at 300g for 2 minutes at 4°C followed by medium removal and resuspension in a mix of buffer Z with enzymes A, P and Y. Single cell suspensions were then blocked with rat anti-mouse CD16/CD32 antibodies (BD Biosciences) used at the recommended dilution for 12 min before incubation with primary antibodies diluted at 1/200 in FACS buffer (PBS, 2% FBS, 0.78 mM EDTA) containing Fc block for 20 min at 4°C. Dead cells were excluded using Live/Dead near-IR dye staining (LIVE/DEAD™ Fixable Near-IR Dead Cell Stain; Invitrogen, L34975) diluted in PBS. PE channel was used for detection of Kaede-Red and Alexa Fluor 488 channel was used for detection of Kaede-Green. 50 μL of CountBright™ Absolute Counting Beads (Invitrogen, C36950) was added to the final sample to measure absolute cell numbers. Flow cytometry data were analysed using FACSDiva software (v4.0) and FlowJo software (Treestar).

### Antibodies

For mouse immunostaining, the following primary antibodies were used: rat anti-mouse B220-AF647 (BioLegend, 103226; clone RA3-6B2, 1/100), rabbit anti-mouse Bassoon (Synaptic System, 141003; 1/200), rabbit anti-mouse c-Fos (Synaptic Systems, 226003; 1/500), rabbit anti-mouse C1q (Abcam, ab182451; 1/200), rat anti-mouse CD11b (Bio-Rad, MCA711G; 1/200), rat anti-mouse CD68 (Bio-Rad, MCA1957; clone FA-11, 1/200), rat anti-mouse CD138 (BD Biosciences, 553712; 1/100),chicken anti-GFP (Abcam, ab13970; 1/500), rabbit anti-mouse GLUT1 (Millipore Sigma, 07-1401; 1/5000), rat anti-HA (Roche, 12158167001; clone 3F10, 1/200), chicken anti-mouse Homer1 (Synaptic Systems, 160006; 1/200), rabbit anti-mouse Iba1 (Wako Chemicals, 019-19741; 1/500), goat anti-mouse IgA (Thermo Fisher Scientific, 62-6700, 1/200), rabbit anti-mouse P2Y12 (Anaspec, AS-55043A; 1/500), rabbit anti-mouse phospho-pyruvate dehydrogenase α1 (Cell Signaling Technology, 37115; 1/500), guinea pig anti-mouse VGLUT1 (Millipore Sigma, AB5905; 1/1000), chicken anti-mouse VGLUT1 (Synaptic Systems, 135316; 1/1000) and guinea pig anti-mouse VGLUT2 (Millipore Sigma, AB2251-I; 1/1000).

Secondary antibodies used were a combination of Alexa Fluor 488, 594 and 647 (1/200, Thermo Fisher Scientific) from either donkey or goat where appropriate. For IgG staining, goat anti-mouse F(ab’)2 AF594 (Thermo Fisher Scientific, A11020) was used.

For flow cytometry, the following antibodies were used: CD16/CD32 (Fc Block; BD Biosciences, clone 2.4G2 (RUO)), CD45-BUV395 (BD Biosciences, clone 30-F11, 563792), CX3CR1-BV421 (BD Biosciences, clone Z8-50, 567531), CD3-BV421 (BD Biosciences, clone 145-2C11, 562600), B220-APC (BD Biosciences, 553092), CD19- BV711 (BioLegend, 115555) and SCA1-PeCy7 (BD Biosciences, 561021).

### Visium spatial transcriptomics

The Visium Gateway Package from 10x Genomics (PN-1000316, 10x Genomics) was used to generate spatial transcriptomic data from two coronal brain sections from two different hM3Dq-expressing CNO-treated mice. Experiments were performed in an RNase-free environment in line with the manufacturer’s recommendations and user guides. The brain was dissected immediately without PBS or PFA perfusion, and fresh frozen directly within an OCT mould in an isopentane bath submerged in liquid nitrogen. The lateral parts of the brain were trimmed during cryosectioning to allow 10 μm-thick coronal sections including both dorsal hippocampi to be cryosectioned into the capture area (6.5 mm x 6.5 mm area) of the Visium Gateway Gene Expression Slide (10x Genomics). Cryosections were taken at 1.9 mm posterior to bregma, more than 2.5 mm anterior to the injection site at 4.6 mm posterior to bregma. Cryosections were also taken adjacent to the sections used for Visium and from MEC to confirm increased ipsilateral GFP and c-Fos immunostaining respectively (data not shown).

The Visium Spatial Tissue Optimization Reagents Kit (10X Genomics) was used to determine an optimal tissue permeabilization time of 12 minutes when performed on 10 μm sections. Samples were mounted directly onto the Visium Gateway Gene Expression Slide and incubated at 37 °C on a thermocycler (T100 Thermal Cycler, Bio-Rad) for one minute, followed by methanol (>99.9%, 34860, Sigma-Aldrich) fixation at −20 °C for 30 minutes. Sections were blocked and stained with DAPI before coverslipping and imaging at 20x magnification using a Zeiss Axioscan 7 microscope. Next, tissue permeabilization was performed as previously to release polyadenylated mRNA for capture by poly(dT) primers on the slide. Steps for reverse transcription, second strand synthesis and denaturation were followed according to the manufacturer’s kit to produce cDNA fragments, from which spatially-barcoded Illumina-compatible libraries were constructed using the dual Index Kit TT Set A (10X Genomics) to add unique i7 and i5 indexes for barcoding.

Single Cell 3’ v3.1 Dual Index libraries were sequenced on a NovaSeq SP instrument by Edinburgh Genomics using parameters recommended by 10X Genomics. These libraries were then demultiplexed by the sequencing provider using the mkfastq command distributed within 10x Genomics’ Cell Ranger software (*65*). The resultant FASTQ was processed in-house by 10x Genomics’ Space Ranger (v1.3.1). The fluorescent microscopy images had to be first manually aligned using 10x Genomics’ Loupe Browser (v6.0.0). Next, ImageJ (v1.53f51) was used to convert images to a TIFF file at the appropriate dimensions. Then within Loupe Browser, an overlay was manually aligned with the fiducial frames present on the Visium slides. Finally, spots for downstream analysis were manually selected, excluding those where tissue was not fully present or was damaged. This produced an alignment file necessary for Space Ranger, which was then run with default parameters. Space Ranger produced a spot-by-count matrix suitable for downstream analysis along with summary statistics confirming successful sequencing and pre-processing. Downstream analysis was carried out in R (v4.0.2) with code available at https://github.com/eturkes/crowley-visium-EC-stim. The Seurat package (v3.2.2) (*66*) was used for all object handling. After reading the count matrix into Seurat, genes completely unexpressed in any spot were excluded. The counts were then normalized using SCTransform (*67*), as recommended by the Seurat authors for handling biologically expected variation in counts due to tissue anatomy. The aim of SCTransform is to maximize biological variation and minimize variation due to technical artifacts by combining information across genes and building a negative binomial distribution model. Both samples were normalized separately for exploratory analysis but normalized together when compared in quantitative analysis.

To identify differentially expressed genes (DEGs) in areas of interest, Loupe Browser was used to manually select spots for comparison between the ipsilateral and contralateral hemispheres. Comparisons included spots in the DGML and the hippocampus overall. Barcodes corresponding to the spots were exported as a CSV file for use in R. After subsetting the data to these spots, SCTransform was applied within the subset, the subset spots were pseudobulked by mouse of origin, and limma (v3.46.0) (*68–70*) was used to identify genes differentially expressed between the ipsilateral and contralateral hemispheres. Limma applies t-statistics in an optimized manner for sequencing data by borrowing information across genes to moderate technical artifacts such as mean-variance trend. The pooling of information across genes also allows null hypothesis with limited sample size, allowing us to use each mouse as true biological replicates rather than a pseudoreplication confounded per-spot analysis. DEGs were defined as those exhibiting changes between comparison groups with an FDR-corrected p-value below 0.05. These genes and others were visualized using functions available within Seurat and Loupe Browser.

### BCR-Seq library preparation and analysis

Animals were deeply anaesthetized and perfused with 20mL ice-cold PBS as previous. Dural meningeal, hippocampal and striatal tissue were collected and immediately placed in RNALater (Invitrogen, AM7020) to preserve RNA integrity. Samples were stored at –80°C until processing. Tissue was homogenized and total RNA extracted using Monarch Total RNA Miniprep Kit (New England Biolabs, T2010S) following manufacturer’s protocol. Reverse transcription and BCR enrichment were performed using NEBNext Immune Sequencing Kit (Mouse) (New England Biolabs, E6330S) in accordance with the manufacturer user manual. BCR-Seq libraries were sequenced by Azenta Lifesciences using a 300bp paired-end sequencing run. The generated BCR-Seq data was processed using MiXCR (v4.7.0) (*71*) using the preset neb-mouse-rna-xcr-umi-nebnext.

Data analysis was performed in the R programming environment (v4.4.1). Clonotype data (considering only heavy chains) from MiXCR were analyzed using the R package immunarch (v0.9.1) (https://CRAN.R-project.org/package=immunarch). Number of unique clonotypes (defined using CDR3 amino acid sequences) for each tissue were extracted, overlaps calculated using the overLapper function in the package systemPipeR (v2.10.0) (*72*) and upset plots generated with UpSetR (v1.4.0) (*73*). For BCR network analysis, the sequencing data were first processed using the nf-core/airrflow (*74*) workflow. The resulting clones file were then filtered to remove non-heavy chains and then passed to dandelion (v0.3.8) (*75*) to generate BCR network plots.

### Confocal imaging

Brain sections for direct comparison were imaged consecutively during the same imaging session. Mounted slides of mouse brain sections were imaged in a single Z-plane using a Plan-Apochromat 10x/0.45 numerical aperture (NA) objective lens initially using the tile function on a Zeiss laser scanning microscope 800 (LSM800) to gain an overview of the mCitrine/EGFP expression and HA immunostaining at the MEC and hippocampal areas. A single-plane image acquired at 10x magnification using the Zeiss LSM800 microscope was also used for imaging of IgA/CD138/IgG immunostaining at the MEC. For c-Fos and pPDH imaging, a Photometrics Prime camera attached to a Zeiss LSM800 microscope was used to acquire widefield images of the MEC region at 10x magnification in 2-3 brain sections per hemisphere. For RNAscope ISH experiments, entire DGML imaging of microglial P2Y12 and *C1qa* mRNA, or entire hippocampal imaging of *Igha*/*Igkc*/*Tnfsf13*/*Tnfrsf17* signal, was performed at 20x magnification, with 3 μm deep Z-stacks (1 μm step size) acquired per brain section and 1-2 sections analysed per hemisphere or per brain for each animal. For measuring IgA and IgG staining at the DG, Z-stack images of 14 μm depth (1 μm step size) per brain section were acquired at 10x magnification and 2 sections analysed per hemisphere. Acquired data was analysed either on ImageJ Fiji or Imaris software where appropriate. Representative images are obtained either from ImageJ Fiji or Imaris and are adjusted for brightness, contrast, and background where appropriate.

### Airyscan confocal imaging and analysis

For imaging of synaptic markers, C1q, VGLUT expression at EGFP+ axon terminals in DGML, and IgG/B220 cells at GLUT1 vessels, the Airyscan function of a Zeiss LSM880 microscope with Plan-Apochromat 63x/1.4 NA oil-immersion objective lens with a zoom factor of 1.8 was used. Optimal frame size and Z-step distance for 3D stacks was determined by applying Nyquist sampling theorem (approximated to 2.3x) to the information frequency to be measured, i.e., axial resolution = ∼400 nm for 488 nm laser => 400/2.3 = 173 nm Z-step size. 8 Z-steps were acquired per stack, meaning a stack thickness of 1.21 μm when 488 nm was the shortest wavelength acquired and 1.47 μm when 594 nm was the shortest wavelength acquired (0.21 μm Z-step size based on Nyquist sampling of 594 nm wavelength). 3 adjacent image stacks (regions of interest; ROIs) were acquired for a given brain region in a given brain section to ensure uniformity of the signal. Deconvolution of Airyscan images was performed using Airyscan processing at strength 6.0 on Zen Black software (v3.0). Deconvoluted output files from Airyscan images were converted to .ims files using Imaris File Converter (v9.5.1). Image files were given pseudonyms to blind from genotype and/or treatment group.

For analysis of synaptic markers and C1q density, .ims image files were loaded into Imaris software (v9.5.1). Channel brightness was adjusted to optimize visualization of immunopositive puncta. The ‘spots’ function was used with ‘Region Growing’, ‘Shortest Distance Calculation’ and ‘Background Subtraction’ functions enabled to select puncta of size 0.2 μm XY diameter and 0.5 μm Z diameter, values based on actual PSF calculations made using sparse fluorescent bead imaging on the Zeiss LSM880 microscope used for these experiments. A threshold was applied to immunoreactive puncta using the ‘Intensity Center’ filter. Threshold values varied between channels and immunostainings but was kept consistent between image pairs being directly compared and between images acquired during the same imaging session. Spot size boundary was defined by ‘Local Contrast’ using a value that appropriately reflected the immunoreactive signal of the source channel. Spots were then filtered for size, where spot volume ≥ 0.007 μm^3^ based on the minimum theoretically resolved spot size using Airyscan of 0.14 μm x 0.14 μm x 0.35 μm = 0.007 μm^3^. For colocalization of Bassoon and Homer1 as a measure of putative excitatory synapse density, a matrix laboratory (MATLAB) script (‘Colocalize Spots’ XTension) was implemented to count presynaptic and postsynaptic spots colocalising with ≤ 250 nm separation between spot centres (*76*). The average value of colocalized spots obtained from a single image was divided by the total image volume to provide an estimate of total synapse density for the ROI. To calculate VGLUT1 and VGLUT2 coverage on GFP+ axons in the DGML, axons were surface rendered based on the GFP channel. Next, VGLUT1 and VGLUT2 puncta were modelled using the spots function as above. The ‘Spots Close to Surface’ XTension was then used with the shortest distance separation set to 0.0 to determine the number of VGLUT1 and VGLUT2 puncta overlapping with the GFP+ axon surface.

### Statistical analysis

Mice included in all analyses were age- and sex-matched littermates wherever possible. WT DREADD surgeries were performed in sets of 4 littermates with EGFP+Veh, EGFP+CNO, hM3Dq+Veh and hM3Dq+CNO treatments performed simultaneously. All analyses were performed blinded to genotype and/or treatment group. Statistical testing was performed on R (v.4.3.2) and GraphPad Prism 10.0 (GraphPad Software). Statistical tests have been performed per experimental replicate, either per animal or per brain hemisphere. Data points shown are animal averages or hemisphere averages (animal/hemisphere average = average of all brain sections, may vary slightly from ROI averages as a result) where points are linked in DREADD experiments. Data shown as mean ± standard error of the mean (SEM) throughout. Statistical tests and data representation are stated in the figure legends. Residuals were tested for normality using a Shapiro-Wilk test and tested for homogeneity of variance by either F test or Levene’s test depending on the data type. Data that failed either test were log_10_ transformed followed by retesting. Where a 2- way mixed ANOVA was used, ‘treatment group’ or ‘brain region’ were used as the between-subjects factor and ‘hemisphere’ was used as the within-subject factor. Significant interaction effects were followed by pairwise t-tests comparing contra- and ipsilateral hemispheres, Bonferroni-adjusting the p-values for multiple testing. Differences between 2 groups were compared using paired or unpaired t-tests as indicated. Throughout, ^ns^p > 0.05, *p < 0.05, **p < 0.01, ***p < 0.001, ****p < 0.0001. Detailed statistical data can be found in Tables S2 and S3.

---

**Figure S1:**
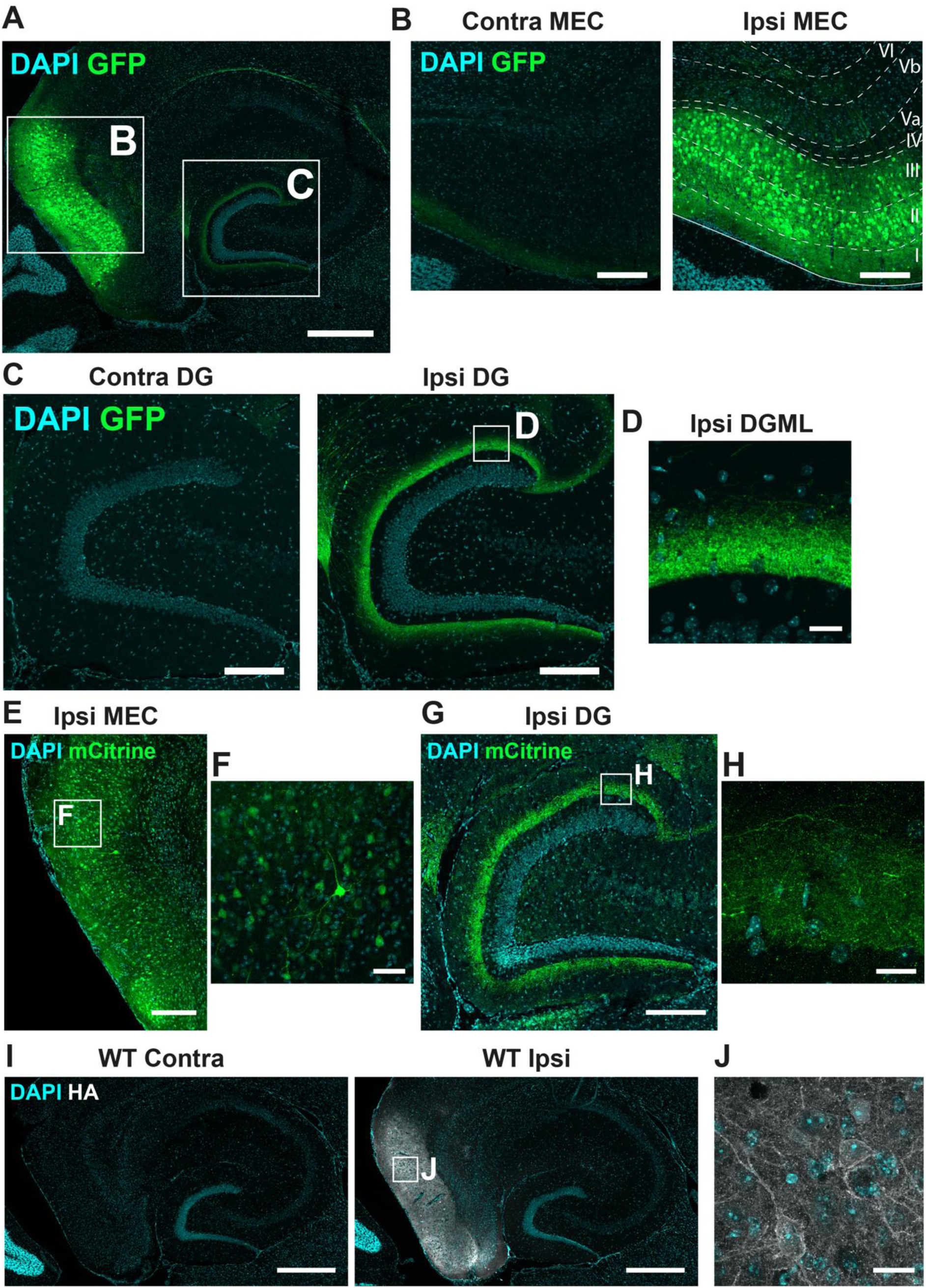
Characterization of MEC viral targeting. **(A)** Representative confocal image of GFP-labeled MEC neurons terminating at the ipsilateral hippocampal DG, scale bar = 250 μm. **(B)** Inset from **(A)** of representative GFP labeling at contralateral and ipsilateral MEC with GFP+ cells restricted to layers II-III; cortical layers delineated in the ipsilateral MEC, scale bar = 100 μm. **(C)** Inset from **(A)** of representative GFP labeling at contralateral and ipsilateral DG with GFP+ projections restricted to the middle part of the DGML, scale bar = 100 μm. **(D)** Inset from **(C)** of GFP+ axonal projections within the middle part of the ipsilateral DGML, scale bar = 25 μm. **(E)** Representative confocal image of mCitrine-labeled MEC neurons (enhanced by anti-GFP antibody) at the ipsilateral MEC, scale bar = 100 μm. **(F)** Inset from **(E)** of representative mCitrine+ cells at the ipsilateral MEC, scale bar = 50 μm. **(G)** Representative mCitrine labeling at ipsilateral DG, scale bar = 100 μm. **(H)** Inset from **(G)** of mCitrine+ projections restricted to the middle part of the DGML, scale bar = 20 μm. **(I)** Absence of HA labeling at the contralateral MEC and presence of HA labeling at the ipsilateral MEC of hM3Dq-expressing CNO-treated mice, scale bar = 250 μm. **(J)** Inset from **(I)** of HA+ cells within the ipsilateral MEC, scale bar = 20 μm.

**Figure S2:**
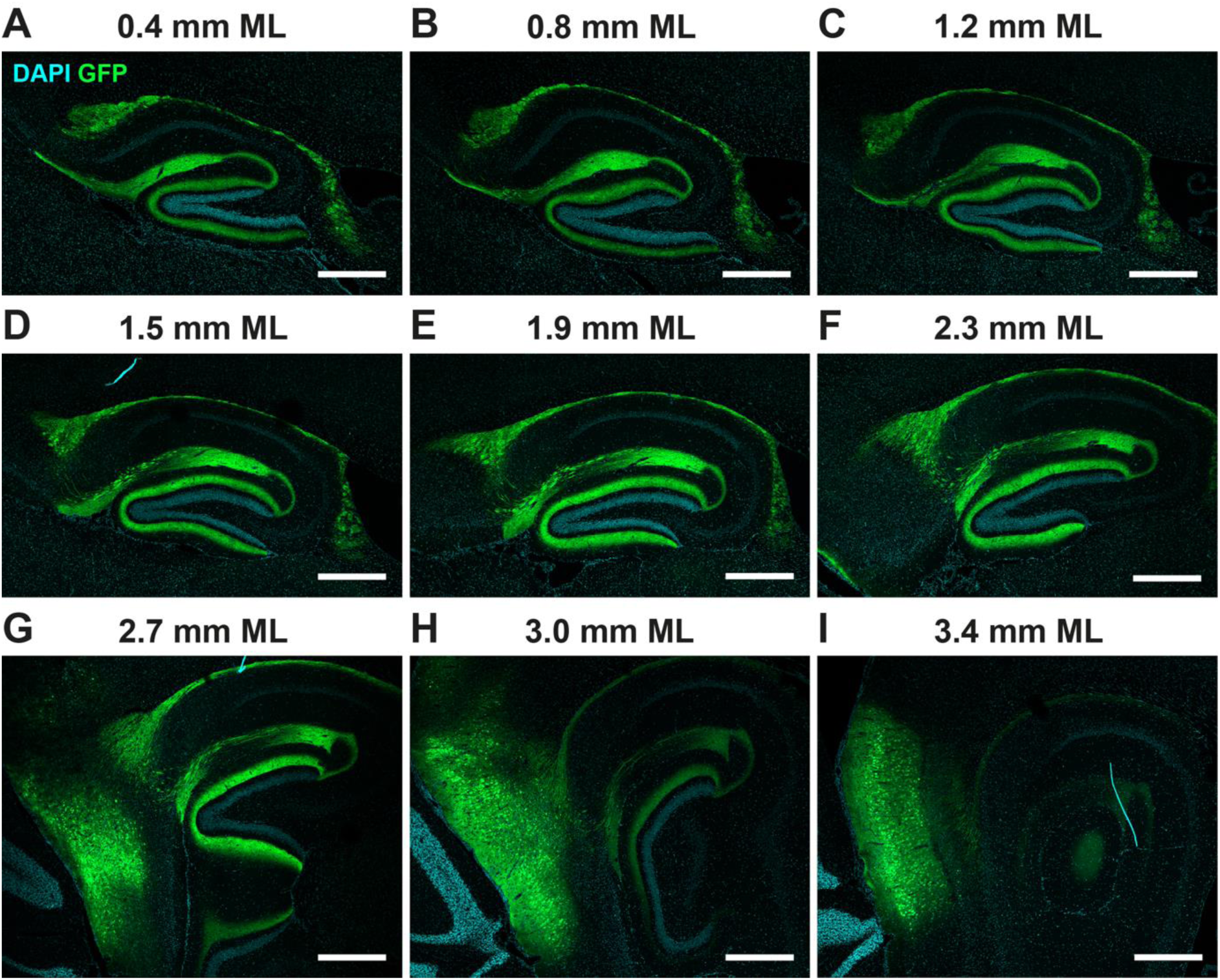
Extent of hippocampal-MEC GFP labeling. **(A)** GFP labeling of sagittal section at 0.4 mm mediolateral. **(B)** 0.8 mm mediolateral. **(C)** 1.2 mm mediolateral. **(D)** 1.5 mm mediolateral. **(E)** 1.9 mm mediolateral. **(F)** 2.3 mm mediolateral. **(G)** 2.7 mm mediolateral. **(H)** 3.0 mm mediolateral. **(I)** 3.4 mm mediolateral, scale bar = 250 μm throughout.

**Figure S3:**
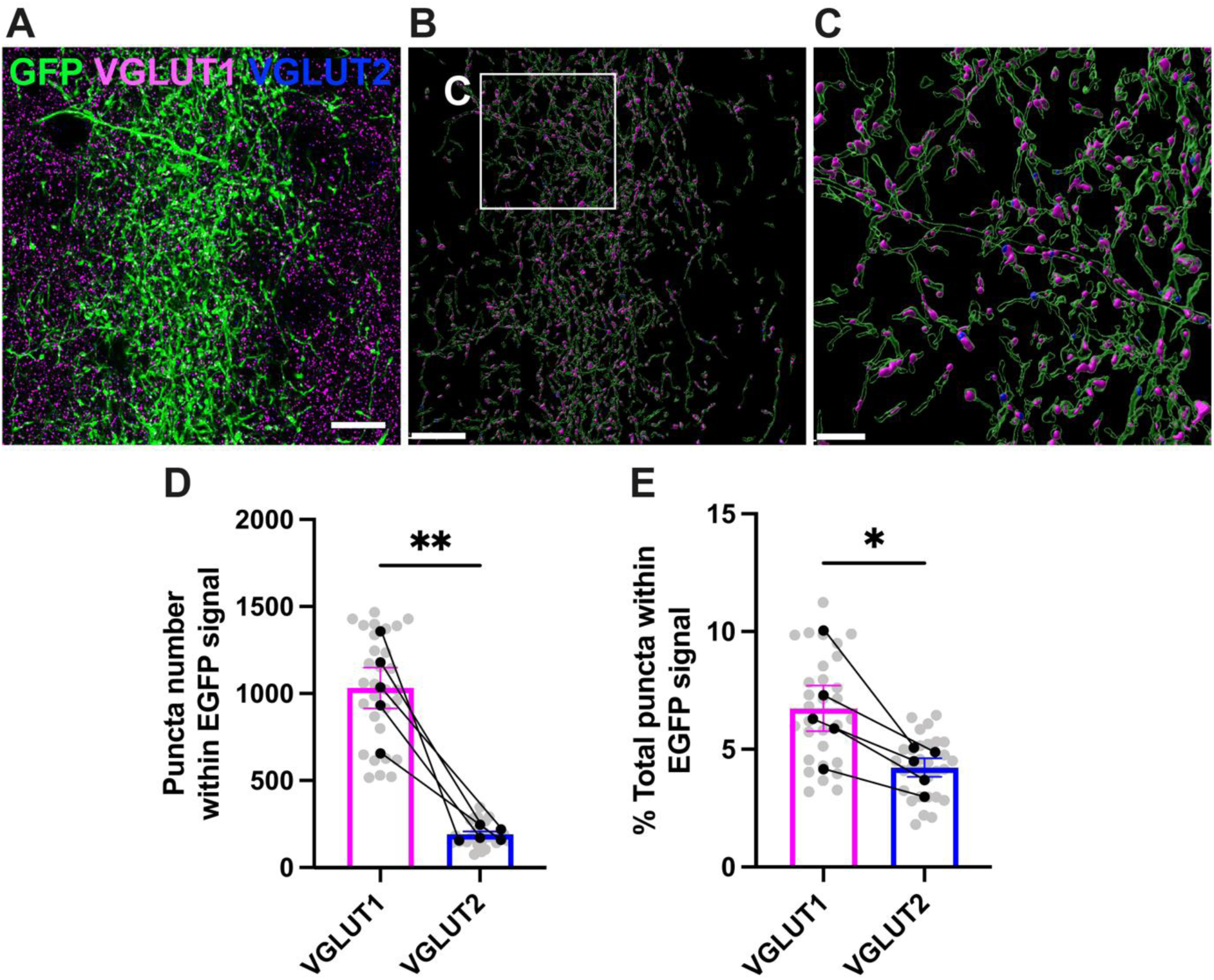
GFP+ terminals contain more VGLUT1 than VGLUT2. **(A)** Representative Airyscan image of GFP projections, VGLUT1 and VGLUT2 presynaptic terminals at the ipsilateral DGML, scale bar = 10μm. **(B)** Imaris 3D reconstruction of **(A)**, scale bar = 10 μm. **(C)** Inset from **(B)** of GFP+ axon projections colocalising with VGLUT1 and VGLUT2, scale bar = 3 μm. **(D)** Quantification of total VGLUT1 puncta and VGLUT2 puncta overlapping with GFP signal at the ipsilateral DGML of hM3Dq+CNO mouse group. 5 EGFP mice (contra = 30 ROIs, ipsi = 30 ROIs), 2 brain sections per mouse, 3 ROIs analysed per section. p-value from paired t test. **(E)** Quantification of percentage of total VGLUT1 puncta and percentage of total VGLUT2 puncta overlapping with GFP signal at the ipsilateral DGML of hM3Dq+CNO mouse group. 5 EGFP mice (contra = 30 ROIs, ipsi = 30 ROIs), 2 brain sections per mouse, 3 ROIs analysed per section. p-value from paired t test. Throughout, circular points represent females, linked points indicate data from the same mouse brain, where points are linked 1 point = 1 hemisphere average = average of brain sections. Data shown as mean ± SEM. ^ns^p > 0.05, *p < 0.05, **p < 0.01.

**Figure S4:**
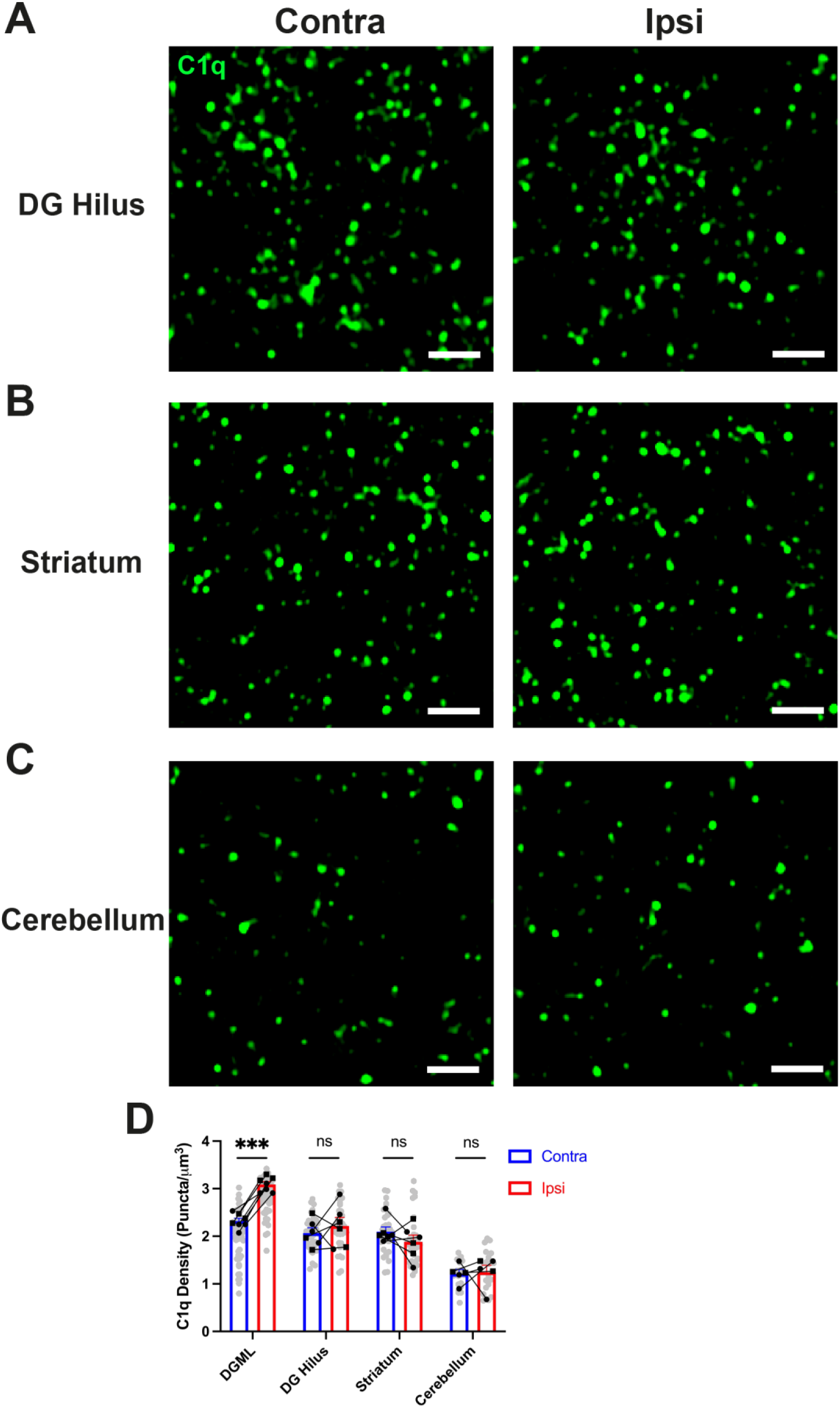
Region-specific activity-dependent C1q upregulation in WT mice. **(A)** Representative Airyscan images of C1q puncta at contralateral and ipsilateral DG hilus of hM3Dq+CNO WT mouse, scale bar = 2 μm. **(B)** Representative Airyscan images of C1q puncta at contralateral and ipsilateral striatum of hM3Dq+CNO WT mouse, scale bar = 2 μm. **(C)** Representative Airyscan images of C1q puncta at contralateral and ipsilateral cerebellum of hM3Dq+CNO WT mouse, scale bar = 2 μm. **(D)** Quantification of C1q puncta density between hemispheres in each brain region. n = 4-7 hM3Dq+CNO mice per region (contra = 54 ROIs, ipsi = 54 ROIs for DGML [identical data to Fig. 2B], contra = 36 ROIs, ipsi = 36 ROIs for DG hilus and striatum, contra = 24 ROIs, ipsi = 24 ROIs for cerebellum), 2 brain sections per hemisphere, 3 ROIs analysed per region. p-values from Bonferroni post-hoc test after 2-way mixed ANOVA with brain region as between- and hemisphere as within-subject factor (interaction p = 0.002). Throughout, square points represent males and circular points represent females, linked points indicate data from the same mouse brain, where points are linked 1 point = 1 hemisphere average = average of brain sections. Data shown as mean ± SEM. ^ns^p > 0.05, ***p < 0.001.

**Figure S5:**
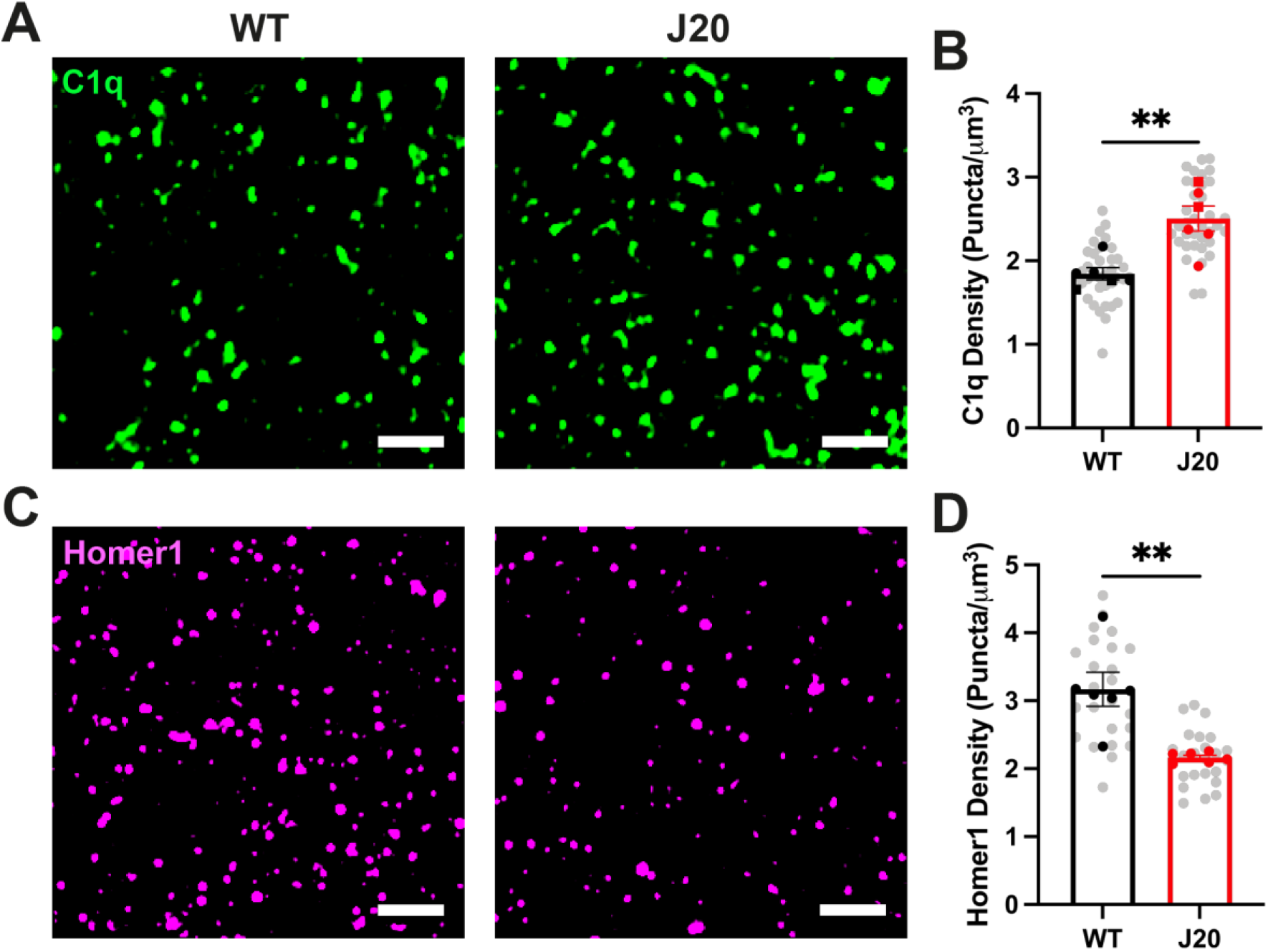
C1q upregulation and Homer1 loss at 4mo in the J20 mouse. **(A)** Representative Airyscan images of C1q puncta at DGML of 4mo WT and J20 mice, scale bar = 2 μm. **(B)** Quantification of C1q puncta density at DGML of 4mo WT and J20 mice. n = 6 mice per group (WT = 36 ROIs, J20 = 36 ROIs), 2 brain sections per mouse, 3 ROIs analysed per section. p-value from unpaired t test. **(C)** Representative Airyscan images of Homer1 puncta at DGML of 4mo WT and J20 mice, scale bar = 2 μm. **(D)** Quantification of Homer1 puncta density at DGML of 4mo WT and J20 mice. n = 6 mice per group (WT = 24 ROIs, J20 = 24 ROIs), 1-2 brain sections per mouse, 3 ROIs analysed per section. p-value from Mann-Whitney U test. Throughout, square points represent males and circular points represent females, 1 point = 1 mouse average = average of brain sections. Data shown as mean ± SEM. ^ns^p > 0.05, **p < 0.01.

**Figure S6:**
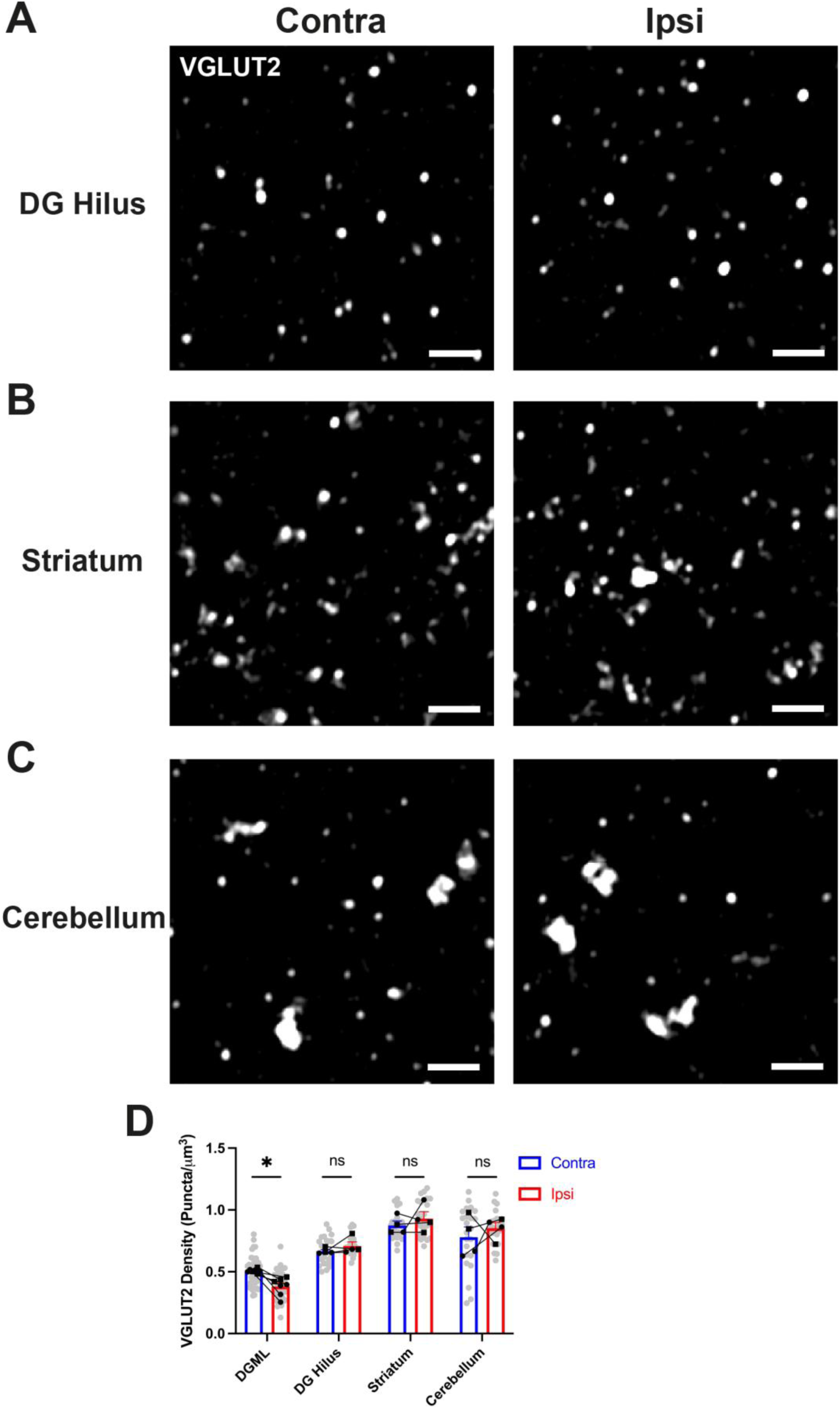
Region-specific activity-dependent VGLUT2 loss in WT mice. **(A)** Representative Airyscan images of VGLUT2 puncta at contralateral and ipsilateral DG hilus of hM3Dq+CNO WT mouse, scale bar = 2 μm. **(B)** Representative Airyscan images of VGLUT2 puncta at contralateral and ipsilateral striatum of hM3Dq+CNO WT mouse, scale bar = 2 μm. **(C)** Representative Airyscan images of VGLUT2 puncta at contralateral and ipsilateral cerebellum of hM3Dq+CNO WT mouse, scale bar = 2 μm. **(D)** Quantification of VGLUT2 puncta density between hemispheres in each brain region. n = 4-7 hM3Dq+CNO mice (contra = 39 ROIs, ipsi = 39 ROIs for DGML [identical data to Fig. 1I], contra = 24 ROIs, ipsi = 24 ROIs for all other regions), 2 brain sections per hemisphere, 3 ROIs analysed per region. p-values from Bonferroni post-hoc test after 2-way mixed ANOVA with brain region as between- and hemisphere as within-subject factor (interaction p = 0.013 after log_10_ transformation). Throughout, square points represent males and circular points represent females, linked points indicate data from the same mouse brain, where points are linked 1 point = 1 hemisphere average = average of brain sections. Data shown as mean ± SEM. ^ns^p > 0.05, *p < 0.05.

**Figure S7:**
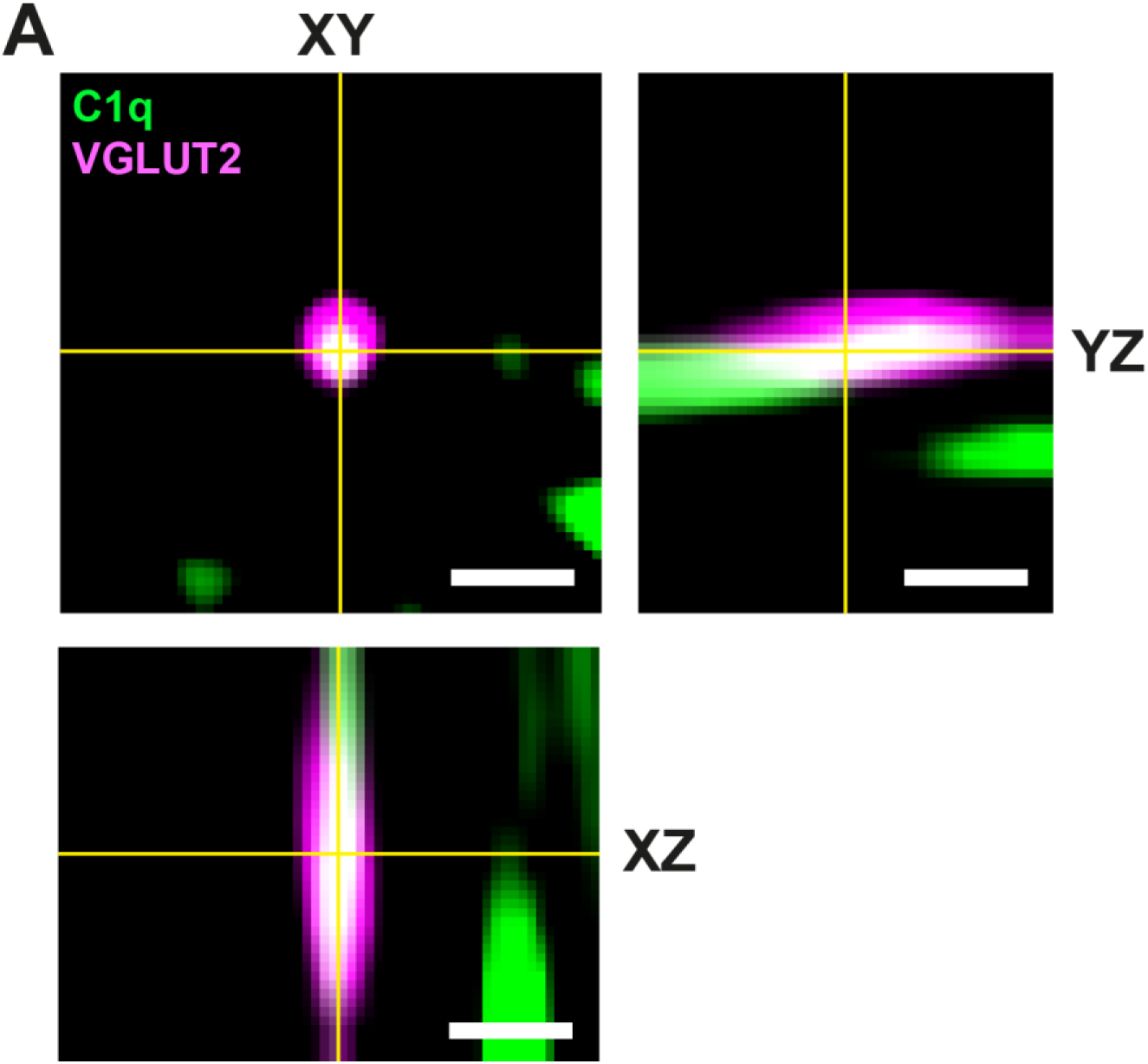
Orthogonal C1q colocalization with VGLUT2 at Airyscan resolution. **(A)** Airyscan resolution representative image and orthogonal views of C1q colocalization with VGLUT2, scale bar = 0.5 μm.

**Figure S8:**
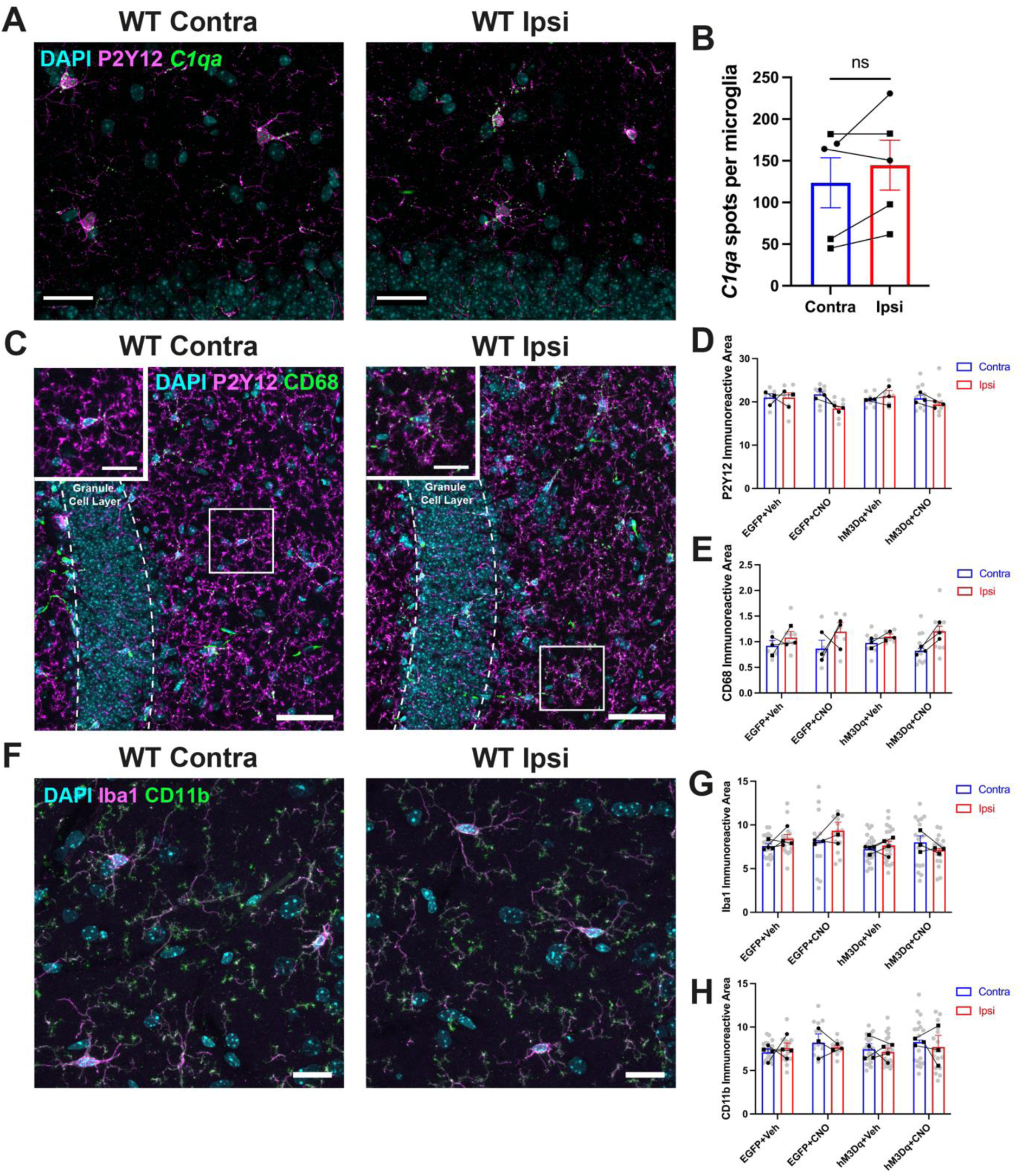
No change in microglial markers at the DGML after hM3Dq expression and/or CNO treatment in perforant pathway projection neurons. **(A)** Representative confocal images of *C1qa* RNAscope and P2Y12 immunostaining at the contralateral and ipsilateral DGML of the hM3Dq+CNO mouse, scale bar = 30 μm. **(B)** Quantification of *C1qa* mRNA puncta per P2Y12+ microglial cell. n = 5 hM3Dq+CNO mice, 1 brain section per hemisphere, 1 ROI analysed per section. p- value from paired t test. **(C)** Representative confocal images of P2Y12 and CD68 at the contralateral and ipsilateral DGML of the hM3Dq+CNO mouse, scale bar = 50 μm. Single DAPI+ microglial cell shown inset, scale bar = 20 μm. **(D)** Quantification of total P2Y12 area at the contralateral and ipsilateral DGML of each mouse group. n = 3 EGFP+Veh mice (contra = 7 ROIs, ipsi = 7 ROIs), n = 3 EGFP+CNO mice (contra = 7 ROIs, ipsi = 7 ROIs), n = 3 hM3Dq+Veh mice (contra = 6 ROIs, ipsi = 6 ROIs) and n = 3 hM3Dq+CNO mice (contra = 10 ROIs, ipsi = 10 ROIs), 2-3 brain sections per hemisphere, 2-3 ROIs analysed per section. Analysed by 2-way mixed ANOVA with group as between- and hemisphere as within-subject factor (interaction p = 0.180). **(E)** Quantification of total CD68 area at the contralateral and ipsilateral DGML of each mouse group. n = 3 EGFP+Veh mice (contra = 7 ROIs, ipsi = 7 ROIs), n = 3 EGFP+CNO mice (contra = 7 ROIs, ipsi = 7 ROIs), n = 3 hM3Dq+Veh mice (contra = 6 ROIs, ipsi = 6 ROIs) and n = 3 hM3Dq+CNO mice (contra = 10 ROIs, ipsi = 10 ROIs), 2 brain sections per hemisphere, 1-3 ROIs analysed per section. Analysed by 2-way mixed ANOVA with group as between- and hemisphere as within-subject factor (interaction p = 0.800). **(F)** Representative confocal images of Iba1 and CD11b at the contralateral and ipsilateral DGML of the hM3Dq+CNO mouse, scale bar = 20 μm. **(G)** Quantification of total Iba1 area at the contralateral and ipsilateral DGML of each mouse group. n = 4 EGFP+Veh mice (contra = 21 ROIs, ipsi = 21 ROIs), n = 3 EGFP+CNO mice (contra = 15 ROIs, ipsi = 15 ROIs), n = 4 hM3Dq+Veh mice (contra = 24 ROIs, ipsi = 24 ROIs) and n = 3 hM3Dq+CNO mice (contra = 18 ROIs, ipsi = 17 ROIs), 2 brain sections per hemisphere, 2-3 ROIs analysed per section. Analysed by 2-way mixed ANOVA with group as between- and hemisphere as within-subject factor (interaction p = 0.265). **(H)** Quantification of total CD11b area at the contralateral and ipsilateral DGML of each mouse group. n = 4 EGFP+Veh mice (contra = 21 ROIs, ipsi = 20 ROIs), n = 3 EGFP+CNO mice (contra = 15 ROIs, ipsi = 15 ROIs), n = 4 hM3Dq+Veh mice (contra = 24 ROIs, ipsi = 24 ROIs) and n = 3 hM3Dq+CNO mice (contra = 18 ROIs, ipsi = 17 ROIs), 2 brain sections per hemisphere, 1-3 ROIs analysed per section. Analysed by 2-way mixed ANOVA with group as between- and hemisphere as within-subject factor (interaction p = 0.847). Throughout, square points represent males and circular points represent females, linked points indicate data from the same mouse brain, where points are linked 1 point = 1 hemisphere average = average of brain sections. Data shown as mean ± SEM. ^ns^p > 0.05.

**Figure S9:**
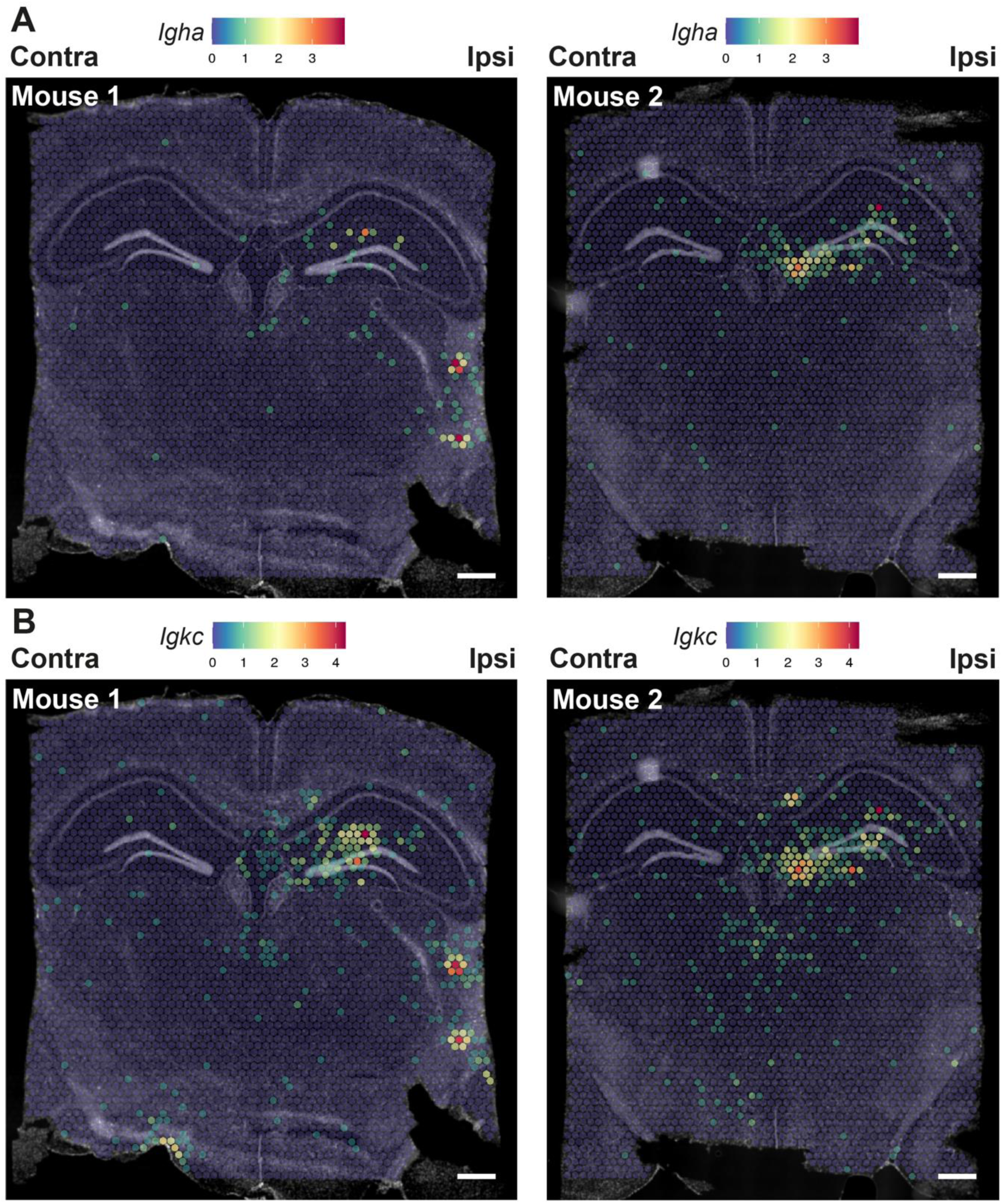
Entire brain section *Igha* and *Igkc* expression using Visium. **(A)** Full brain section images of Visium spot mRNA transcript counts for *Igha*, scale bar = 500 μm. **(B)** Full brain section images of Visium spot mRNA transcript counts for *Igkc*, scale bar = 500 μm.

**Figure S10:**
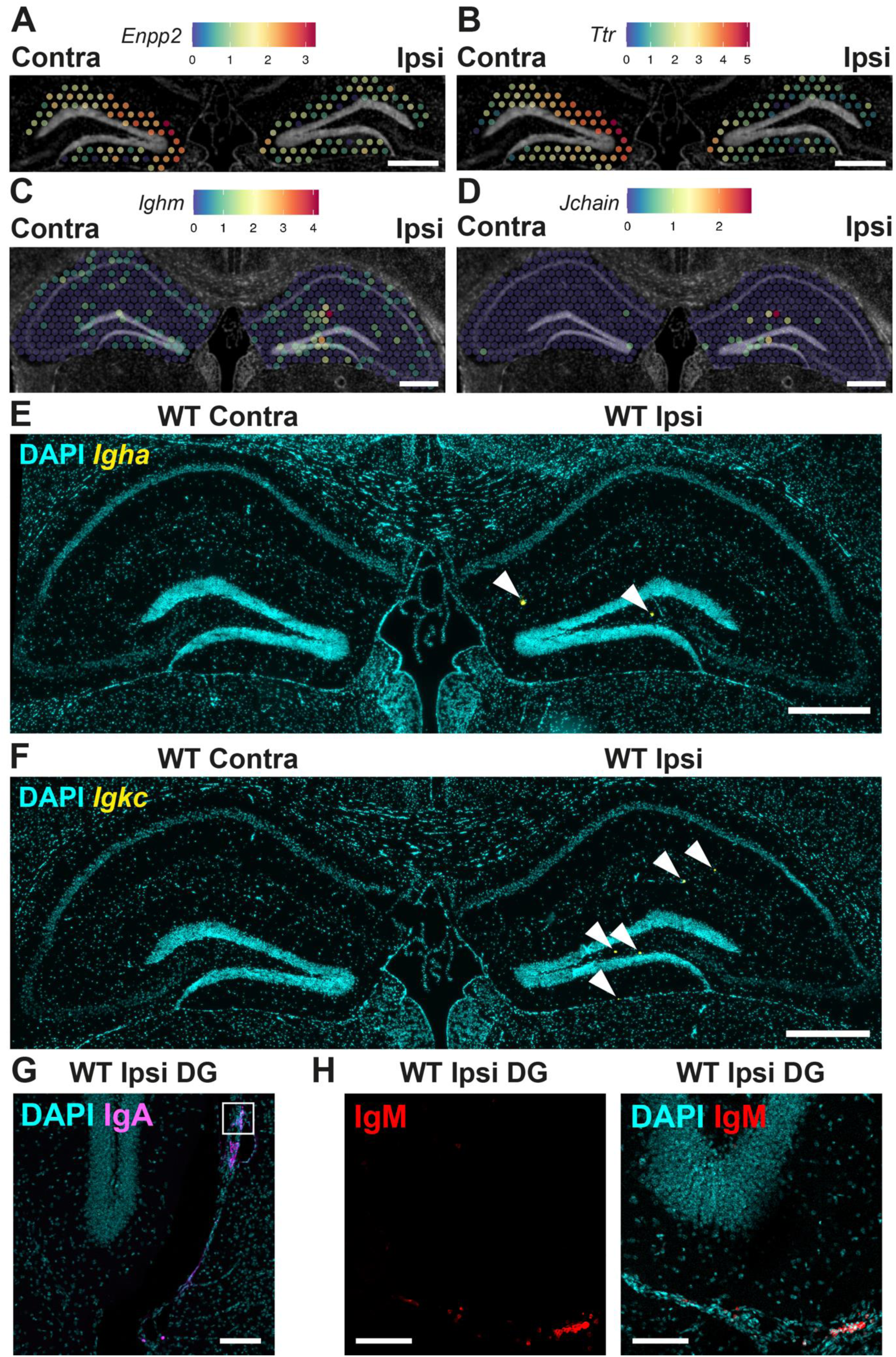
Activity-dependent Visium changes and immunoglobulins. **(A)** Representative images of Visium spot mRNA transcript counts at contralateral and ipsilateral DGML for *Enpp2*, scale bar = 500 μm. **(B)** Representative images of Visium spot mRNA transcript counts at contralateral and ipsilateral DGML for *Ttr*, scale bar = 500 μm. **(C)** Representative images of Visium spot mRNA transcript counts at contralateral and ipsilateral DGML for *Ighm*, scale bar = 500 μm. **(D)** Representative images of Visium spot mRNA transcript counts at contralateral and ipsilateral DGML for *Jchain*, scale bar = 500 μm. **(E)** Representative confocal image of *Igha* RNAscope at contralateral and ipsilateral hippocampi, scale bar = 500 μm. White arrowheads indicate *Igha*+ cells. **(F)** Representative confocal image of *Igkc* RNAscope at contralateral and ipsilateral hippocampi, scale bar = 500 μm. White arrowheads indicate *Igkc*+ cells. **(G)** Representative confocal image of IgA at ipsilateral DG, scale bar = 100 μm. Inset represents IgA+ CD138+ cells in Fig. 3D. **(H)** Representative confocal image of IgM at ipsilateral DG, scale bar = 100 μm.

**Figure S11:**
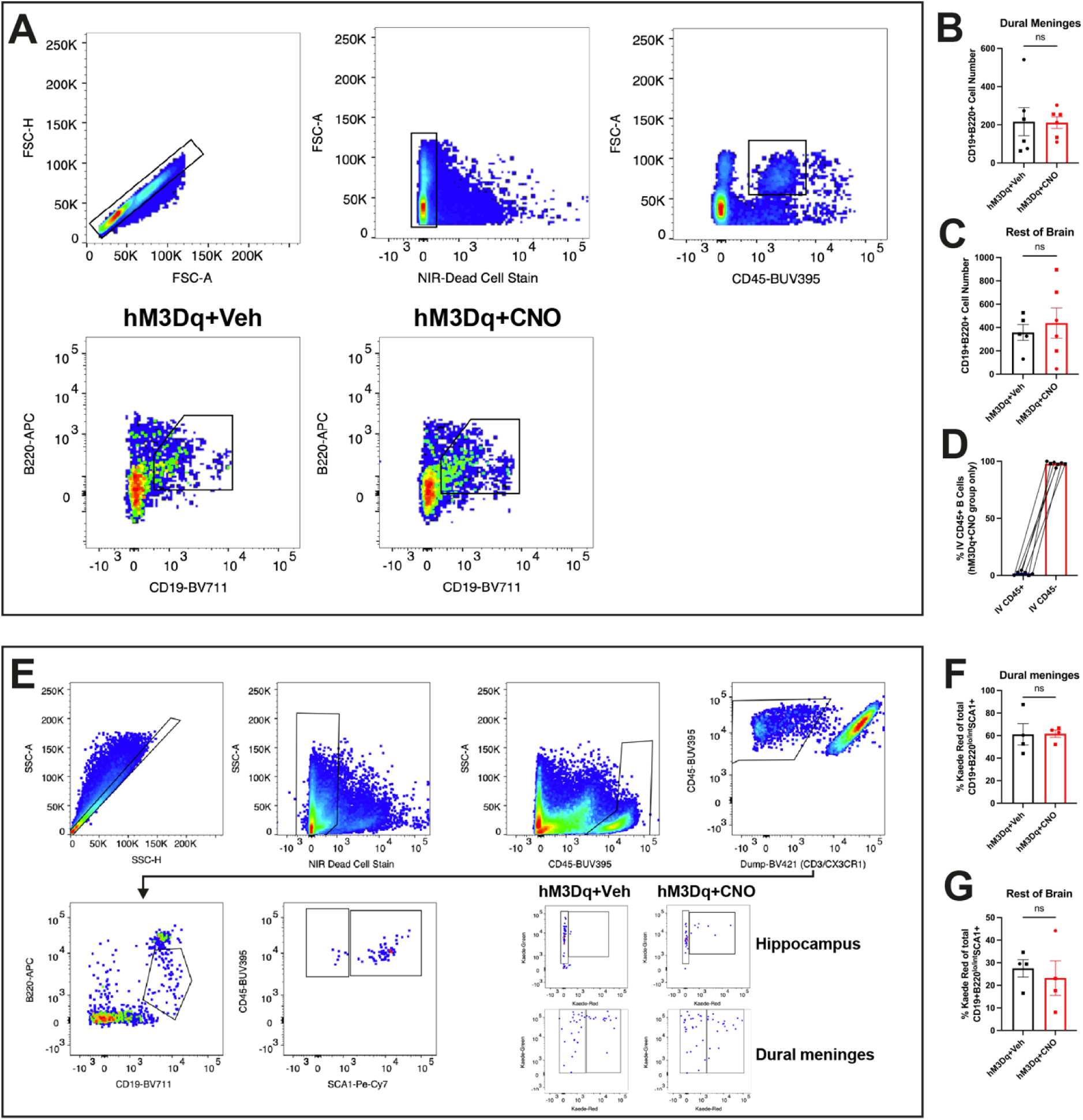
FACS analysis of B lymphocyte lineage cells. **(A)** FACS gating strategy to quantify CD19+B220+ cells from mouse brain tissue and representative FACS plots from hM3Dq+Veh and hM3Dq+CNO mice. **(B)** Quantification of CD19+B220+ cells in dural meninges of n = 6 hM3Dq+Veh and n = 6 hM3Dq+CNO mice. p-value from unpaired t test. **(C)** Quantification of CD19+B220+ cells in brain tissue without the hippocampus (denoted ‘rest of brain’) of n = 5 hM3Dq+Veh and n = 6 hM3Dq+CNO mice. p-value from unpaired t test. **(D)** Quantification of percentage hippocampal CD19+B220+ cells from n = 7 hM3Dq+CNO mice labeled with intravenous CD45 antibody. **(E)** FACS gating strategy to quantify CD19+B220^Lo/Int^SCA1+ cells from mouse brain tissue. **(F)** Quantification of % Kaede Red CD19+B220^Lo/Int^SCA1+ cells in dural meninges of n = 4 hM3Dq+Veh and n = 4 hM3Dq+CNO mice. p-value from unpaired t test. **(G)** Quantification of % Kaede Red CD19+B220^Lo/Int^SCA1+ cells in non-hippocampal brain tissue (denoted ‘rest of brain’) of n = 4 hM3Dq+Veh and n = 4 hM3Dq+CNO mice. p-value from unpaired t test. Throughout, square points represent males and circular points represent females. Data shown as mean ± SEM. ^ns^p > 0.05, ****p < 0.0001.

**Figure S12:**
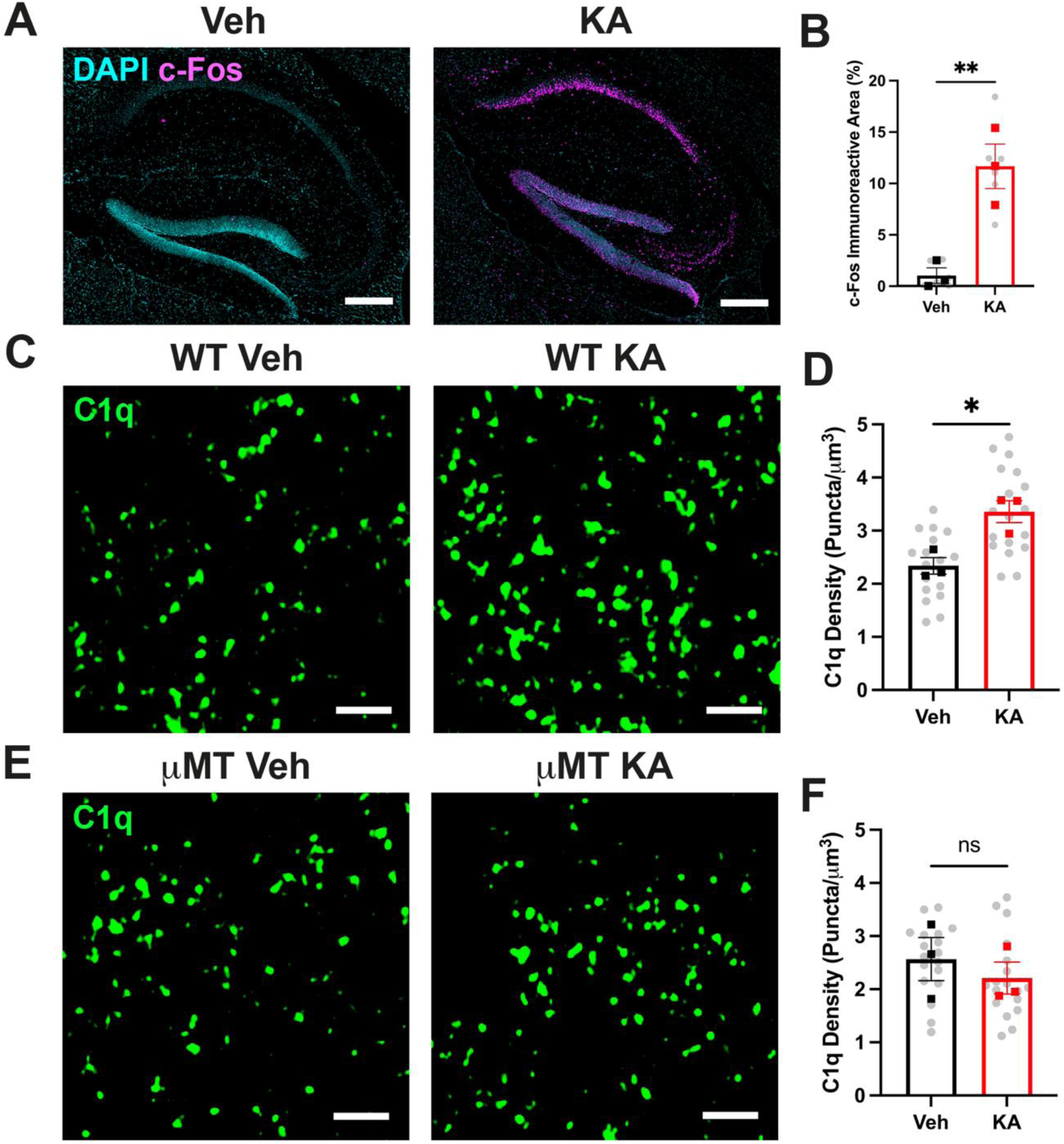
C1q upregulation upon KA treatment requires *Ighm* expression. **(A)** Representative confocal images of c-Fos immunostaining at the hippocampus of mice treated with vehicle or KA, scale bar = 250 μm. **(B)** Quantification of c-Fos immunopositive area between DG after vehicle and KA treatment. n = 3 WT mice per group, 2 brain sections per mouse, 1 ROI analysed per section. p-value from unpaired t test. **(C)** Representative Airyscan images of C1q puncta at CA3SL of mice treated with vehicle or KA, scale bar = 2 μm. **(D)** Quantification of C1q puncta density after vehicle and KA treatment. n = 3 WT mice per group (veh = 18 ROIs, KA = 18 ROIs), 2 brain sections per mouse, 3 ROIs analysed per section. p-value from unpaired t test. **(E)** Representative Airyscan images of C1q at CA3SL of vehicle-treated and KA- treated μMT mice, scale bar = 2 μm. **(F)** Quantification of C1q puncta density at CA3SL of vehicle-treated and KA-treated μMT mice. n = 3 μMT mice per group (veh = 18 ROIs, KA = 18 ROIs), 2 brain sections per mouse, 3 ROIs analysed per section. p-value from unpaired t test. Throughout, square points represent males, 1 point = 1 mouse = average of brain sections. Data shown as mean ± SEM. ^ns^p > 0.05, *p < 0.05, **p < 0.01.

**Table S1:** Modified Racine scale. Adapted from (*64*).

| Stage Number | Description |
| --- | --- |
| 1 | Freezing behaviour |
| 2 | Rigid posture with raised tail |
| 3 | Head bobbing and forepaw shaking |
| 4 | Rearing, falling and jumping |
| 5 | Continuous level 4 |
| 6 | Loss of posture and generalized convulsions |

**Table S2:**
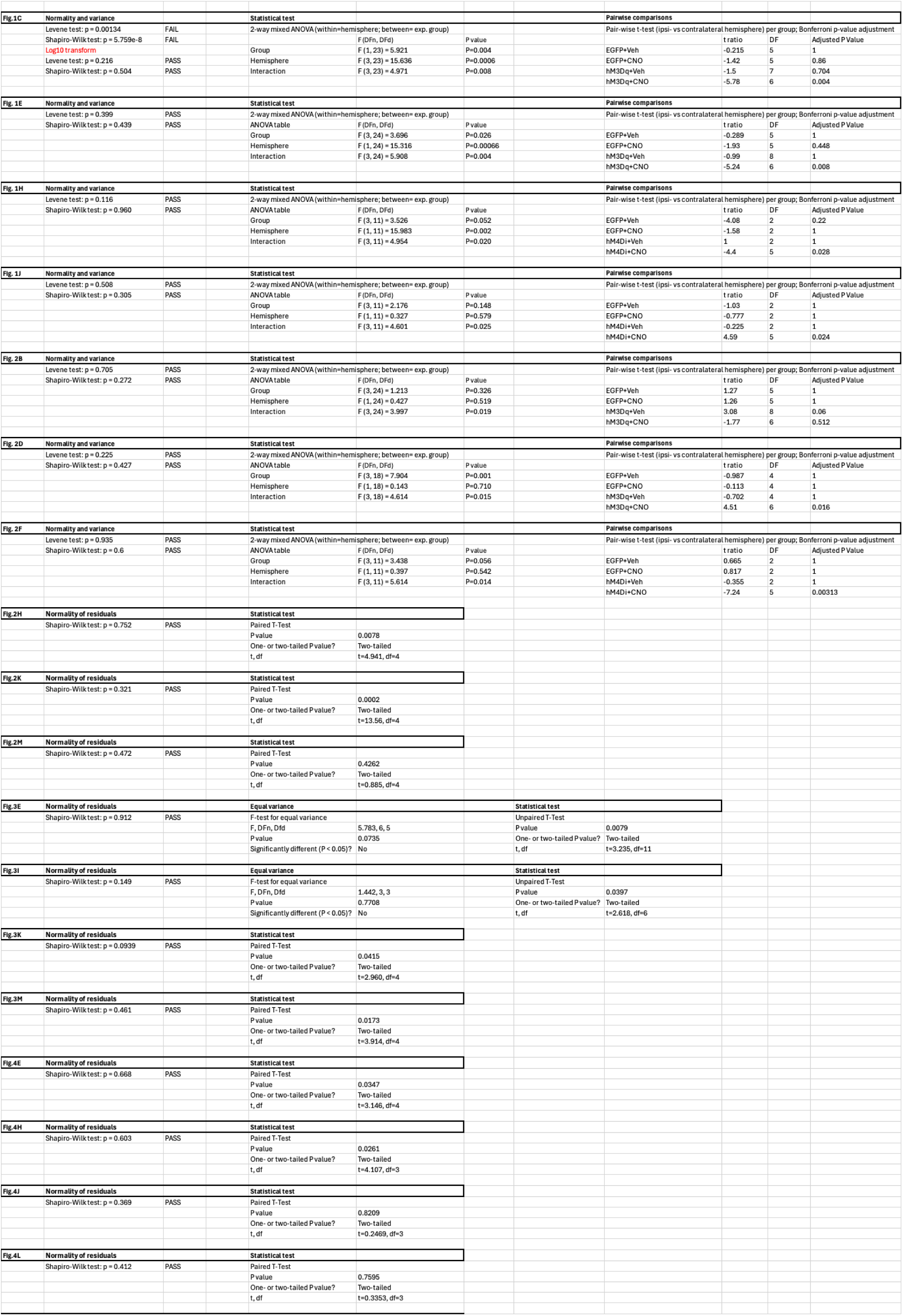
Detailed statistical data from main figures.

**Table S3:**
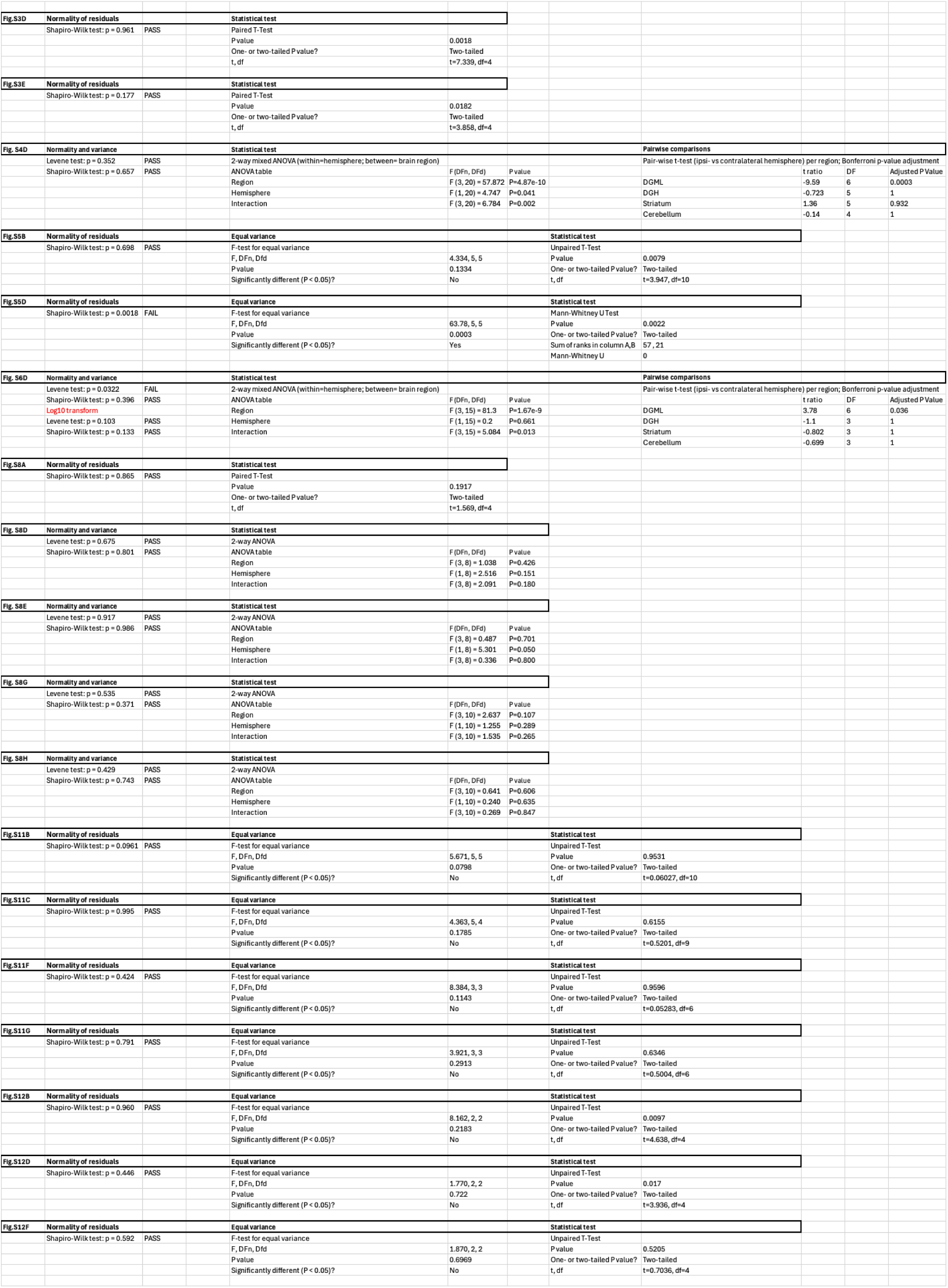
Detailed statistical data from supplementary figures.

